# Timing the origin of eukaryotic cellular complexity with ancient duplications

**DOI:** 10.1101/823484

**Authors:** Julian Vosseberg, Jolien J. E. van Hooff, Marina Marcet-Houben, Anne van Vlimmeren, Leny M. van Wijk, Toni Gabaldón, Berend Snel

**Affiliations:** Theoretical Biology and Bioinformatics, Department of Biology, Faculty of Science, Utrecht University, Utrecht, The Netherlands; Centre for Genomic Regulation, The Barcelona Institute of Science and Technology, Barcelona, Spain; Life Sciences Department, Barcelona Supercomputing Center, Barcelona, Spain; Mechanisms of Disease, Institute for Research in Biomedicine, Barcelona, Spain; Institució Catalana de Recerca i Estudis Avançats, Barcelona, Spain

## Abstract

Eukaryogenesis is one of the most enigmatic evolutionary transitions, during which simple prokaryotic cells gave rise to complex eukaryotic cells. While evolutionary intermediates are lacking, gene duplications allow us to elucidate the order of events by which eukaryotes originated. Here we use a phylogenomics approach to reconstruct successive steps during eukaryogenesis. We found that gene duplications roughly doubled the proto-eukaryotic genome, with families inherited from the Asgard archaea-related host being duplicated most. By relatively timing events using phylogenetic distances we inferred that duplications in cytoskeletal and membrane trafficking families were among the earliest events, whereas most other families expanded primarily after mitochondrial endosymbiosis. Altogether, we demonstrate that the host that engulfed the proto-mitochondrion had some eukaryote-like complexity, which further increased drastically upon mitochondrial acquisition. This scenario bridges the signs of complexity observed in Asgard archaeal genomes to the proposed role of mitochondria in triggering eukaryogenesis.

## Introduction

Compared to prokaryotes, eukaryotic cells are tremendously complex. Eukaryotic cells are larger, contain more genetic material, have multiple membrane-bound compartments and operate a dynamic cytoskeleton. The last eukaryotic common ancestor (LECA) already had an intracellular organisation and gene repertoire characteristic of present-day eukaryotes^1^, making the transition from particular prokaryotes to the first eukaryotes – eukaryogenesis – one of the main unresolved puzzles in evolutionary biology^2,3^.

Most eukaryogenesis models involve a host, related to the recently discovered Asgard archaea^4,5^, which took up an Alphaproteobacteria-related endosymbiont^6,7^ that gave rise to the mitochondrion. However, the timing and impact of this endosymbiosis event in the evolution of eukaryotic complexity are hotly debated and at the heart of different scenarios on eukaryogenesis^8^.

Besides the acquisition of genes via the endosymbiont, the proto-eukaryotic genome expanded through gene inventions, duplications and horizontal gene transfers during eukaryogenesis^9,10^. Previous work suggested that gene duplications nearly doubled the ancestral proto-eukaryotic genome^10^. Gene families such as small GTPases, kinesins and vesicle coat proteins greatly expanded, which enabled proto-eukaryotes to employ an elaborate intracellular signalling network, a vesicular trafficking system and a dynamic cytoskeleton^11–14^.

Uncovering the order in which these and other eukaryotic features emerged is complicated due to the absence of intermediate life forms. However, duplications occurred during the transition and are likely to yield valuable insights into the intermediate steps of eukaryogenesis. In this study we attempt to reconstruct the successive stages of eukaryogenesis by systematically analysing large sets of phylogenetic trees. We determined the scale of gene inventions and duplications and how different functions and phylogenetic origins had contributed to these eukaryotic innovations. Furthermore, we timed the prokaryotic donations and duplications relative to each other using information from phylogenetic branch lengths.

## Results

### Unprecedented resolution of duplications during eukaryogenesis

To obtain a comprehensive picture of duplications during eukaryogenesis we made use of the Pfam database (see Methods)^15^. Instead of selecting a few eukaryotic species beforehand we chose to take an approach inspired by the ‘ScrollSaw’ method^13^, which limits phylogenetic analyses to slowly evolving sequences and collapses duplications after LECA. We constructed phylogenetic trees and detected 10,259 nodes in these trees that were inferred to represent a Pfam domain in LECA (‘LECA families’) (Fig. 1a). To include genes having only small Pfam domains, which were excluded for computational reasons, or no domains at all, we used a linear regression analysis to obtain an estimated LECA genome containing 12,780 genes (95% prediction interval: 7,463 – 21,886) (Extended Data Fig. 1).

**Fig. 1.**
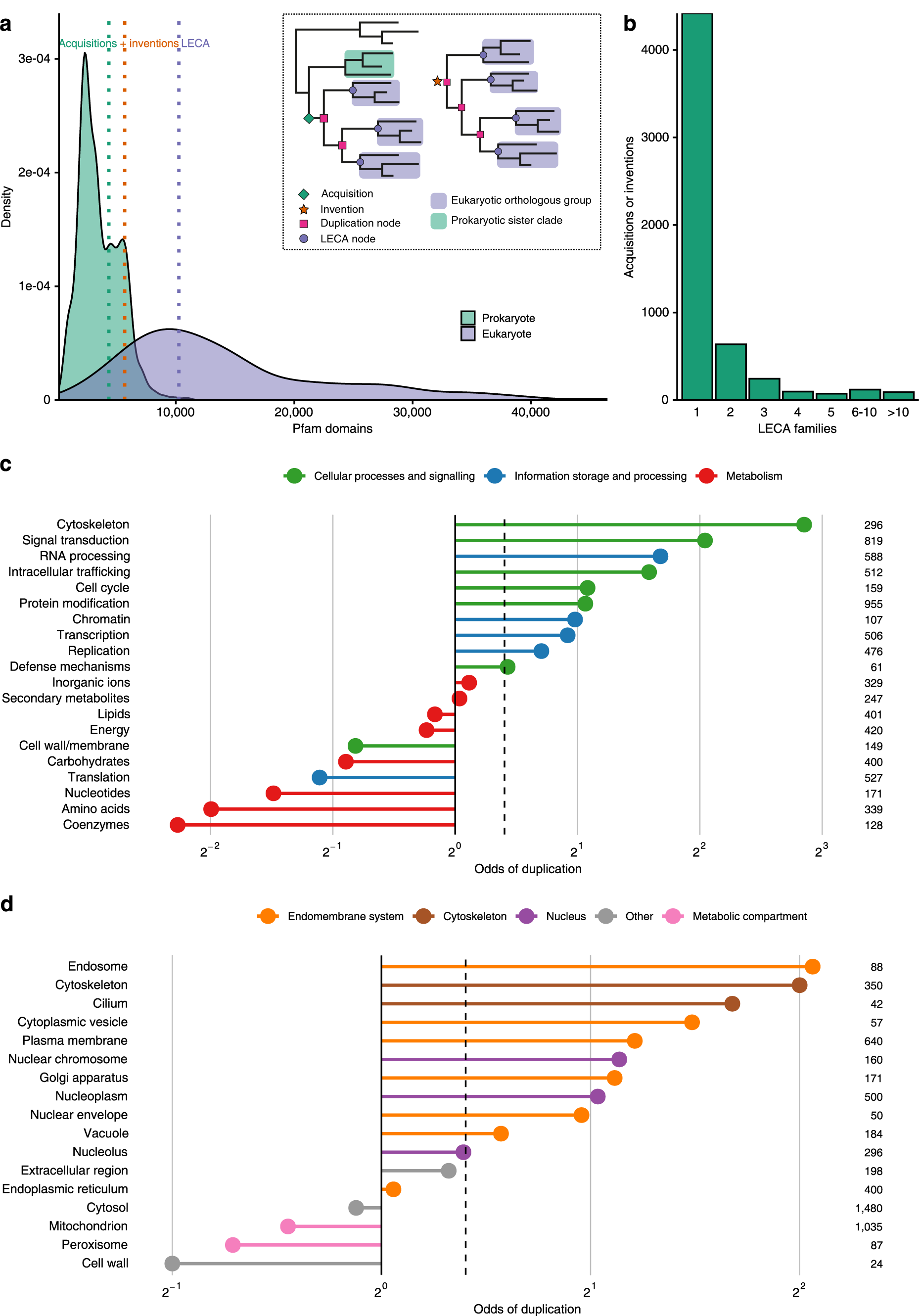
Characterisation of duplications during eukaryogenesis. **a**, Density plot showing the distribution of the number of Pfam domains in present-day prokaryotes (green) and eukaryotes (purple) in comparison with the acquisition, invention and LECA estimates obtained from phylogenetic trees (see inset). **b**, Number of acquisitions or inventions that gave rise to a particular number of LECA families, demonstrating the skewedness of duplications across protein families. **c**, Odds of duplication for LECA families according to KOG functional categories. 78% of pairwise comparisons were significantly different (Supplementary Fig. 1). The poorly characterised categories and functions of very few families (cell motility, extracellular structures and nuclear structure) are not depicted. **d**, Odds of duplication for LECA families according to cellular localisation. 60% of pairwise comparisons were significantly different (Supplementary Fig. 2). **c-d**, Numbers indicate the number of LECA families and dashed lines indicate the odds or fraction of all LECA families in total.

Comparing the number of inferred LECA families to extant eukaryotes showed that the genome size of LECA reflected that of a typical present-day eukaryote (Fig. 1a), which is in in line with the inferred complexity of LECA, but in contrast with lower estimates obtained previously^10,16^. The multiplication factor – the number of LECA families divided by the number of acquired and invented genes or domains – was 1.8, approximating the near doubling reported before^10^. The observed doubling was validated in an additional data set (Extended Data Table 1), despite a recent study that inferred very few duplications during eukaryogenesis (see Supplementary Information)^17^. Although on average genes duplicated once, the distribution of duplications is heavily skewed with many acquisitions or inventions not having undergone any duplication (Fig. 1b). The enormous expansion of the proto-eukaryotic genome was dominated by massive duplications in a small set of families (Extended Data Table 2).

There is a considerable difference between duplicated and non-duplicated LECA families regarding their functions and cellular localisations. Metabolic LECA families rarely had a duplication history, whereas LECA families involved in information storage and processing, and cellular processes and signalling were more likely to descend from a duplication (χ^2^ = 619, df = 2, P = 3.4 × 10^−135^; Fig. 1c, Supplementary Fig. 1). Notable exceptions to this pattern were families involved in cell wall or membrane biogenesis and translation, which were rarely duplicated. The observed differences in functions were reflected by the differences between cellular localisations, with proteins in the endomembrane system and cytoskeleton mostly resulting from a duplication (χ^2^ = 288, df = 4, P = 4.1 × 10^−61^; Fig. 1d, Supplementary Fig. 2). Like duplications, inventions primarily occurred to families involved in informational and cellular processes (χ^2^ = 228, df = 2, P = 3.7 × 10^−50^ (function); χ^2^ = 180, df = 4, P = 5.3 × 10^−38^ (localisation); Supplementary Fig. 3-6). For complex eukaryotes to emerge, most innovations occurred in nuclear processes, the endomembrane system, intracellular transport and signal transduction, especially due to gene duplications.

**Fig. 2.**
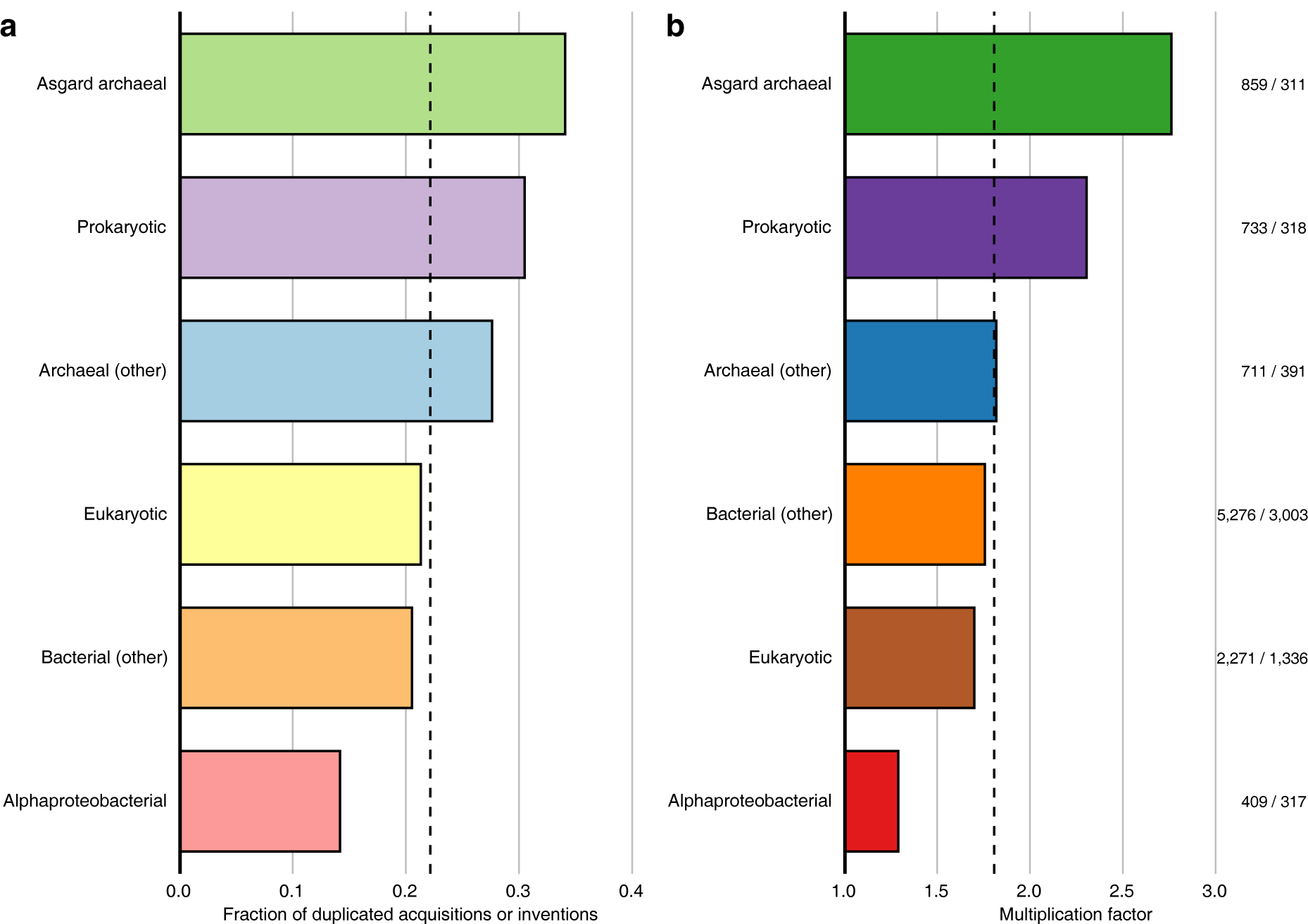
Contribution of different phylogenetic origins to duplications during eukaryogenesis. **a**, Duplication tendency as fraction of clades having undergone at least one duplication. **b**, Multiplication factors, defined as the number of LECA families divided by the number of acquisitions or inventions. These numbers are shown beside the corresponding bar. **a, b**, Dashed lines indicate the duplication tendency and multiplication factor for all acquisitions and LECA families. Prokaryotic: unclear prokaryotic ancestry (could not be assigned to a domain or lower taxonomic level).

**Fig. 3.**
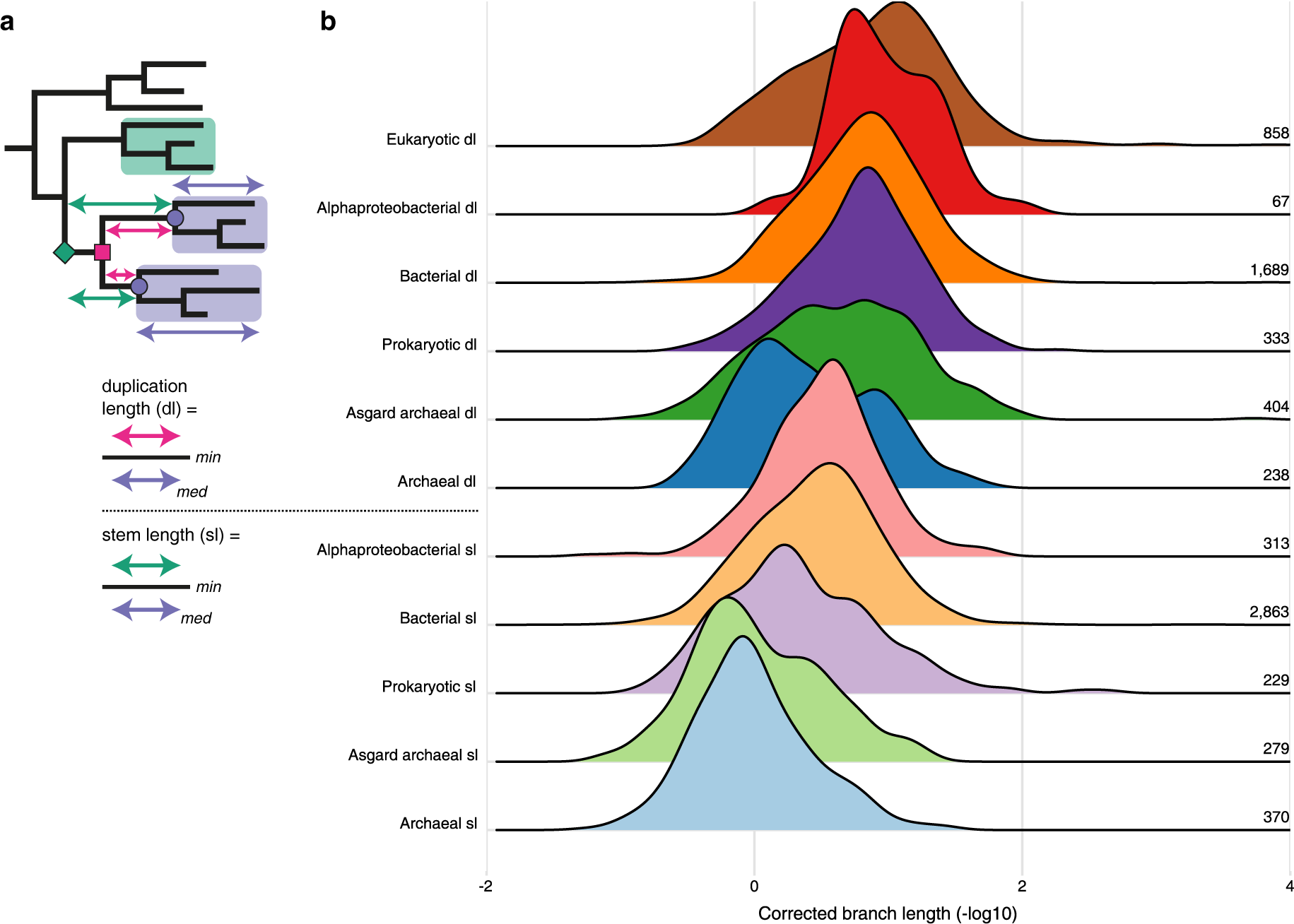
Timing of acquisitions and duplications from different phylogenetic origins during eukaryogenesis. **a**, Illustration how the stem and duplication lengths were calculated. Symbols and colour schemes are identical to Fig. 1a. The phylogenetic distance between the acquisition or duplication and LECA were normalised by dividing it by the median branch length between LECA and the eukaryotic terminal nodes. In case of duplications the shortest of the possible normalised paths was used. **b**, Ridgeline plot showing the distribution of corrected stem or duplication lengths, depicted as the additive inverse of the log-transformed values. Consequently, longer branches have a smaller value and vice versa. For clarity, a peak of near-zero branch lengths is not shown (see Extended Data Fig. 4). Numbers indicate the number of acquisitions or duplications for which the branch lengths were included. Groups are ordered based on the median value.

### Relatively large contribution of the host to duplicated LECA families

For the Pfams that were donated to the eukaryotic stem lineage we identified the prokaryotic sister group, which represents the best candidate for the Pfam’s phylogenetic origin (Extended Data Fig. 3a). Most acquisitions had a bacterial sister group (76%), of which only a small proportion was alphaproteobacterial (7% of all acquisitions), in agreement with previous analyses^9,18,19^. The acquisitions from archaea (16%) predominantly had an Asgard archaeal sister (9% of all acquisitions). Moreover, the most common Asgard archaeal sister group was solely comprised of Heimdallarchaeota (Extended Data Fig. 3b); especially Heimdallarchaeote LC3 was frequently the sister group. This is in line with previous analyses providing support for either all Heimdallarchaeota or LC3 being the currently known archaeal lineage most closely related to eukaryotes^20,21^. The species in alphaproteobacterial sister groups, on the other hand, came from different orders (Extended Data Fig. 3c), consistent with the recently proposed deep phylogenetic position of mitochondria^7^. The remaining acquisitions (7%) had an unclear prokaryotic ancestry (see Supplementary Information).

Families with different sister clades varied substantially in the number of gene duplications they experienced during eukaryogenesis (χ^2^ = 62, df = 5, P = 4.8 × 10^−12^ (duplication tendency); χ^2^ = 293, df = 5, P = 2.7 × 10^−61^ (LECA families from duplication); Fig. 2, Supplementary Fig. 7). The multiplication factor of 2.8 for families likely inherited from the Asgard archaea-related host was strikingly high compared with the other groups (between 1.3 and 2.3). Especially duplications related to the ubiquitin system and trafficking machinery contributed to the relatively large number of host-related paralogues (Extended Data Table 2). In contrast, there was a clear deficit of duplications in families with an alphaproteobacterial sister group (multiplication factor of 1.3). Hence, the endosymbiont marginally contributed to the near doubling of the genetic material during eukaryogenesis, whereas the host contributed relatively the most.

### Branch lengths point to a mitochondria-intermediate scenario

The striking differences in duplication dynamics between families with different affiliations could tentatively stem from differences in timing of these acquisitions and subsequent duplications. For example, the low number of alphaproteobacterial-associated duplications could be the result of a late mitochondrial acquisition. Branch lengths have previously been used to time the acquisition of genes from the different prokaryotic donors^9^. Duplications were not included in that analysis. Although the measure has been criticised for its assumption that rates pre- and post-LECA are correlated^22,23^, it did yield correct timings for certain post-LECA events^9,24^. Shorter branch lengths, corrected for differences in evolutionary rates across families, reflect more recent acquisitions. Similarly, duplications can be timed with the length of the branch connecting the duplication and LECA nodes (Fig. 3a). Using branch lengths to time duplications is a novel implementation of this approach, which we here use to infer the order of events during eukaryogenesis.

For the timing of acquisitions we obtained similar results as before^9^, with archaeal stems being longer than bacterial stems (P = 4.5 × 10^−63^, two-sided Mann-Whitney *U*-test; Fig. 3b, Supplementary Fig. 8). Among the archaeal stem lengths the Asgard archaeal stems were shortest, as were the alphaproteobacterial stems among the bacterial stems, although these differences failed to reach statistical significance in both cases (P = 0.31 and P = 0.07, respectively). Figure 3b shows that there is a wide distribution of host-related duplication lengths, with a substantial number of duplication lengths both longer and shorter than (alphaproteo)bacterial stem lengths. This pattern is independent of the normalisation (Extended Data Fig. 4). Endosymbiont-related and invented families showed the shortest duplication lengths. These differences in branch lengths indicate that an increase in genomic complexity via duplications had already occurred prior to the mitochondrial acquisition.

**Fig. 4.**
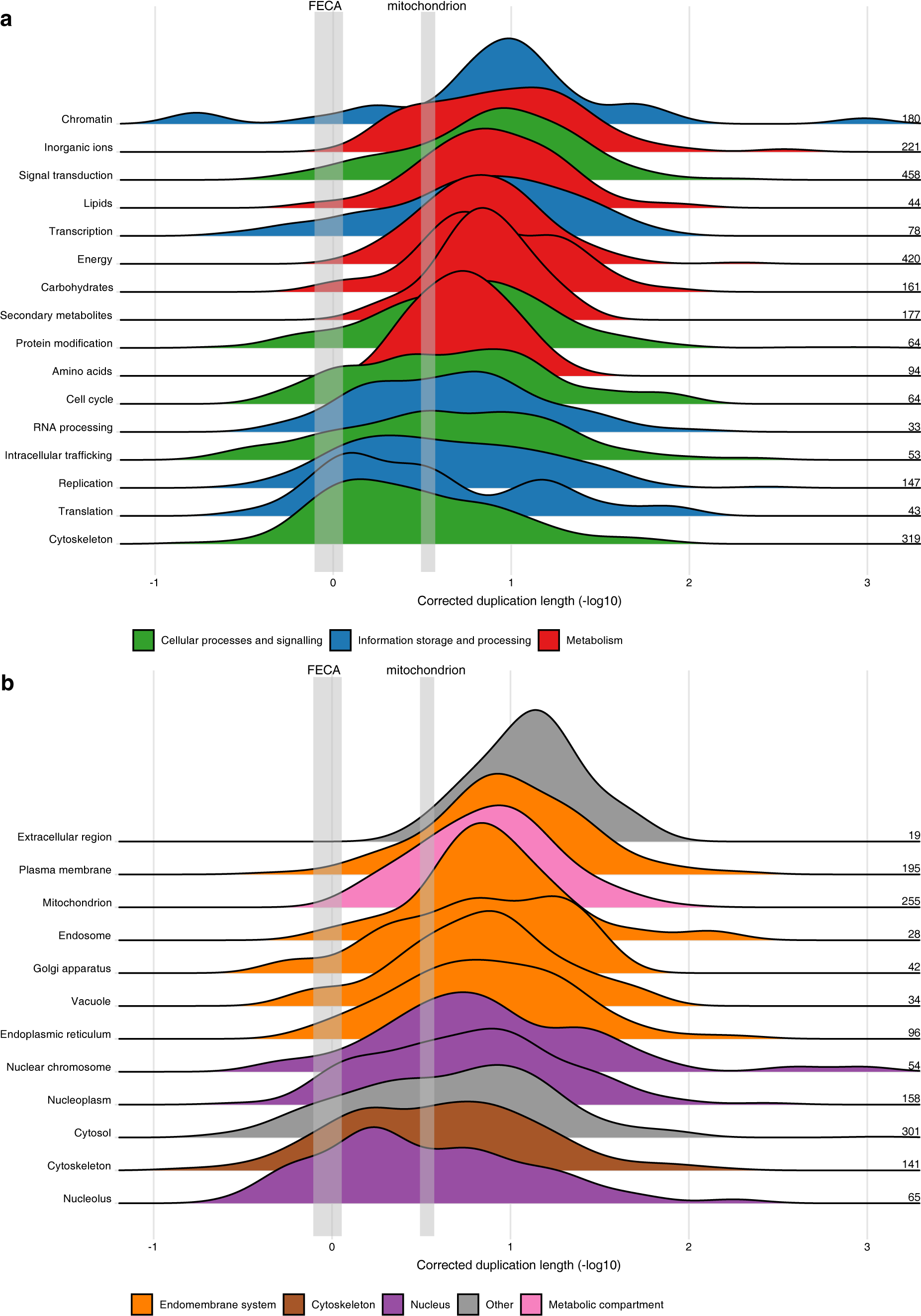
Timing of duplications during eukaryogenesis according to function and localisation. **a, b**, Ridgeline plots showing the distribution of duplication lengths for different functional categories (**a**) and cellular localisations (**b**). The binomial-based 95% confidence interval of the median of the Asgard archaeal (FECA) and alphaproteobacterial stem lengths (mitochondrion) are depicted in grey, indicating the divergence of eukaryotes from their Asgard archaea-related and Alphaproteobacteria-related ancestors, respectively. Groups are ordered based on the median value.

To make inferences about cellular complexity we categorised the duplications according to their functional annotations and cellular localisations. A marked distinction in duplication lengths between different functions can be observed, with duplications in metabolic functions corresponding to shorter branches (P = 6.6 × 10^−7^, Kruskal-Wallis test; Fig. 4a, Supplementary Fig. 9). Moreover, a substantial number of duplication lengths in information storage and processes, and cellular processes and signalling were longer than the alphaproteobacterial stem length, including multiple corresponding to duplications assigned to the cytoskeleton and intracellular trafficking. Duplications in signal transduction and transcription families mainly had shorter branch lengths, indicating that these regulatory functions evolved relatively late. With respect to cellular localisation, nucleolar and cytoskeletal duplication lengths were longest and most duplications related to the endomembrane system had duplication lengths shorter than the alphaproteobacterial stem length (Fig. 4b, Supplementary Fig. 10), indicating that the increase in cellular complexity before the mitochondrial acquisition mainly comprised the evolution of cytoskeletal and nucleolar components.

## Discussion

This large-scale analysis of duplications during eukaryogenesis provides compelling evidence for a mitochondria-intermediate eukaryogenesis scenario. The results suggest that the Asgard archaea-related host already had some eukaryote-like cellular complexity, such as a dynamic cytoskeleton and membrane trafficking. Upon mitochondrial acquisition there was an even further increase in complexity with the establishment of a complex signalling and transcription regulation network and by shaping the endomembrane system. These post-endosymbiosis innovations could have been facilitated by the excess of energy allegedly provided by the mitochondrion^25,26^.

A relatively complex host is in line with the presence of homologues of genes functioning in the eukaryotic cytoskeleton and membrane trafficking in Asgard archaeal genomes^4,5,27^. Moreover, some of them, including ESCRT-III homologues, small GTPases and (loki)actins, have duplicated in these archaea as well, either before eukaryogenesis or more recently^4,5,27^. This indicates that there has already been a tendency for at least the cytoskeleton and membrane remodelling to become more complex in Asgard archaeal lineages. A dynamic cytoskeleton and trafficking system, perhaps enabling primitive phagocytosis^28^, might have been essential for the host to take up the bacterial symbiont. Molecular and cell biology research in these archaea, from which the first results have recently become public^29,30^, is highly promising to obtain more insight into the nature of the host lineage. In addition to a reconstruction of the host, further exploration of the numerous acquisitions, inventions and duplications during eukaryogenesis is key to fully unravelling the origin of eukaryotes.

## Acknowledgements

We thank K. S. Marakova and E. V. Koonin for sharing their KOG-to-COG protein clusters with us. We are grateful to T. J. P. van Dam, E. S. Deutekom and G. J. P. L. Kops for useful advice and discussions. This work is part of the research programme VICI with project number 016.160.638, which is (partly) financed by the Netherlands Organisation for Scientific Research (NWO). T.G. acknowledges support from the Spanish Ministry of Science and Innovation for grant PGC2018-099921-B-I00 and from the European Union’s Horizon 2020 research and innovation programme under grant agreement ERC-2016-724173.

## Author contributions

J.J.E.v.H., T.G. and B.S. conceived the study. J.V. and J.J.E.v.H. performed the research. J.V., J.J.E.v.H., T.G. and B.S. analysed and interpreted the results. M.M.H. and A.v.V. aided in the development of the tree analysis pipeline. L.M.v.W. implemented the ScrollSaw-based method. J.V., J.J.E.v.H. and B.S. wrote the manuscript, which was edited and approved by all authors.

## Competing interests

The authors declare no competing interests.

## Methods

### Data

The 209 eukaryotic (predicted) proteomes were from an in-house dataset that has been used and described before^31^. Prokaryotic proteomes (3,457 in total) were extracted from eggnog 4.5^32^. The prokaryotic dataset was supplemented with nine predicted proteomes from the recently described Asgard superphylum^5^.

### Pfam analysis

#### Pfam assignment

We used hmmsearch (HMMER v3.1b2^33^) with the Pfam 31.0 profile hidden Markov models (HMMs)^15^ and the corresponding gathering thresholds to assess to which Pfam what part of each prokaryotic and eukaryotic sequence should be assigned. The domains that were hit were extracted from the sequences based on the envelope coordinates. If a sequence had hits to multiple Pfams and these hits were overlapping for at least 15 amino acids only the best hit was used. If the same Pfam had multiple hits in the sequence due to an insertion relative to the model the different hits were artificially merged. Since the latter is more prone to errors for short models and short sequences contain less phylogenetic signal, profile HMMs shorter than fifty amino acids were not considered for further analysis.

#### Reduction of sequences

For each Pfam, the number of prokaryotic sequences was reduced with kClust v1.0^34^ using a clustering threshold of 2.93. Asgard archaeal sequences were excluded from this reduction, because they are relatively undersampled and are already genetically diverse.

The number of eukaryotic sequences was reduced with a novel method (L.M.v.W. and B.S., manuscript in preparation) based on the ScrollSaw approach^13^. For each Pfam an all species versus all species BLAST^35^ was performed. Because we were only interested in the best hit the max_target_seqs option was set to 1. Although this option has raised some attention recently^36,37^, we only used it as a proxy for evolutionary distance and our analysis would not be seriously impacted by this option given the overall small sizes of our databases. Subsequently, bidirectional best hits (BBHs) between sequences from different supergroups (Excavata, Archaeplastida + Cryptista, SAR + Haptista, Obazoa and Amoebozoa) or between Diphoda (first three supergroups) and Opimoda (other two supergroups), which we consider a likely root of the eukaryotic tree of life^38^, were identified. The corresponding sequences were used for phylogenetic analysis.

#### Phylogenetic analysis

Multiple sequence alignments were made with MAFFT v7.310^39^ (auto option) and trimmed with trimAl v1.4.rev15^40^ (gap threshold 10%). Phylogenetic trees were inferred with IQ-TREE v1.6.4^41^ (LG4X model^42^, 1000 ultrafast bootstraps^43^). If the consensus tree had a higher likelihood than the best tree from the search, the first was used for further analysis. Because inferring trees for PF00005 (ABC transporters) in this way was too computationally demanding, we used FastTree v2.1.10^44^ with the LG model to construct trees for this Pfam. This Pfam was not considered for branch length analysis (see below).

### KOG-COG clusters analysis

#### Selecting sequences

Clusters of homologous sequences were created based on the KOG-to-COG mappings established by Makarova *et al*.^10^. This dataset contains eukaryotic orthologous groups (KOGs) mapped to homologous prokaryotic orthologous groups (COGs). In many cases, multiple KOGs had been mapped to a single COG, which often reflects a duplication. Furthermore, KOGs had been clustered if they are homologous to each other but lack a homologous COG.

To assign sequences to the KOG-to-COG clusters, we applied a specific method for each of the following three species categories: the eukaryotes, the prokaryotes excluding Asgard archaea, and the Asgard archaea. For eukaryotes, we collected protein sequences from 10 species present in 5 eukaryotic supergroups to obtain a good representation of eukaryotic diversity: *Naegleria gruberi* and *Euglena gracilis* (Excavata), *Cladospihon okarmurans* and *Bigelowiella natans* (SAR+Haptista), *Guillardia theta* and *Klebsormidium flaccidum* (Archaeplastida+Cryptista), *Acanthamoeboa castellanii* and *Acytostelium subglobosum* (Amoebozoa), and *Capsaspora owczarzaki* and *Nuclearia* sp. (Obazoa). These species were selected because they were most often in BBHs in the Pfam sequence selection (see above). We derived profile HMMs for euNOGs from eggNOG 4.5^32^. The original KOG-to-COG clusters also contained ‘TWOGs’, candidate orthologous groups. For each TWOG we found the best matching ‘ENOG’ (‘Unsupervised Cluster of Orthologous Group’) provided by eggNOG. We combined the profile HMMs of these ENOGs with the KOG profile HMMs and created a profile database. We performed hmmscan (HMMER v3.1b1^33^) to assign protein sequences from the eukaryotic species to KOGs/ENOGs. For the COGs present in the KOG-to-COG clusters, we obtained all their member sequences. We downloaded profile HMMs of all COGs from eggNOG and assigned the Asgard protein sequences to COGs using hmmscan. Subsequently, for all KOGs (and ENOGs, which we collectively refer to as KOGs) and COGs, we reduced the number of sequences with kClust v1.0^34^, using a score per column of 3.53. We subsequently merged homologous sequences from eukaryotes, prokaryotes and Asgard archaea according to the KOG-to-COG mapping, resulting in novel KOG-to-COG clusters.

#### Phylogenetic analyses

For each KOG-to-COG cluster, we generated phylogenetic trees using an in-house pipeline also used previously^9^. The sequences were aligned using MAFFT v6.861b^45^, option –auto, and subsequently trimmed using trimAl v1.4^40^ with a gap threshold of 0.1. From these alignments, we constructed phylogenetic trees using FastTree v2.1.8^44^ with WAG as evolutionary model.

### Tree analyses

#### Removal of interspersing prokaryotes

Trees were analysed with an in-house ETE3^46^ script. We examined whether the tree contained prokaryotic sequences that probably reflect recent horizontal gene transfers and that might interfere with our analysis. Prokaryotic sequences from a single genus that were in between eukaryotic sequences were pruned from the tree. If there was only one prokaryotic sequence it was kept only if it was an Asgard archaeal sequence, because it has been reported that sometimes only a single sequenced Asgard archaeon contains a sequence otherwise only present in eukaryotes^5^.

#### Annotation of eukaryotic nodes

For each eukaryotic clade the nodes were annotated as duplications prior to LECA, LECA nodes, post-LECA nodes or unclassified. Only clades that contained at least one LECA node were of interest. The node combining the eukaryotic clade with the rest of the tree (if present) was annotated as acquisition node.

For the annotation of nodes in trees from the Pfam-ScrollSaw sequences the information from the eukaryotic sequences that were not in the between-supergroup or Opimoda-Diphoda BBHs were included. To correctly assign in-paralogues we additionally performed an own species versus own species BLAST for each Pfam (max_target_seqs 2). The sequences belonging to a Pfam that were not in the tree were mapped onto their best hits in the tree according to the BLAST score.

In order to infer reliable duplication nodes in the tree, duplication consistency scores were calculated for all internal nodes starting from the root of a eukaryotic clade. This score is the overlap of species at both sides of a node divided by the total number of species at both sides, taking both sequences in the tree and assigned sequences (as described above) into account. If the duplication consistency score was at least 0.2 and both daughter nodes fulfilled the LECA criteria, this node was annotated as duplication node. A LECA node first had to have both Opimoda and Diphoda sequences in the clade. Besides, to take care of post-LECA horizontal gene transfer (HGT) events among eukaryotes and of tree uncertainties, the mean presence of a potential LECA family in eukaryotic species, weighted so that each supergroup contributes equally, should be at least 15%. If a node did not fulfil the LECA criteria it was annotated as a post-LECA node.

The abovementioned thresholds were chosen based on manual inspection of a selection of trees. Using different thresholds for duplication consistency (0, 0.1, 0.2, 0.3) and LECA coverage scores (0, 5, 10, 15, 20 and 25%) did not have a large impact on the absolute numbers and quality measures, such as the fraction of well-supported LECA and duplication nodes (data not shown). This underlines the robustness of our analysis.

For the annotation of nodes in KOG-COG trees a slightly different approach was followed. Because of the lower number of eukaryotic species a species overlap criterion of 2 was used for duplication nodes instead of consistency scores. If both Opimoda and Diphoda sequences were among the descendants of a node, it fulfilled the LECA criterion.

After this first annotation round all LECA nodes were re-evaluated. If there were duplication nodes in both daughters, this node had to be a duplication node as well even though its duplication consistency score was below the threshold. If there were duplication nodes in only one daughter lineage, the LECA node was annotated as unclassified. It could reflect a duplication event or a tree artefact due to rogue taxa. If there were no duplication nodes in either daughter lineages, all LECA nodes in the daughter lineages of this LECA node were reannotated as post-LECA nodes.

#### Rooting eukaryote-only trees

For trees with only eukaryotic sequences and trees for which all prokaryotic sequences had been removed, inferring the root poses a challenge. For these trees duplication and LECA nodes were called in unrooted mode. The distances between the LECA nodes were calculated and the tree was rooted in the middle of the LECA nodes that were furthest apart, resulting in an additional duplication node at this root. If there were no duplications found in this way, because there were less than two duplications in the tree, rooting was tried on each internal node. The node that fulfilled the duplication criteria and that maximised the species overlap was chosen. If none fulfilled the criteria, it was checked if the entire tree fulfilled the LECA criteria. For Pfams for which we could not infer a tree because there were only two or three sequences selected, we also checked if this Pfam in itself fulfilled the LECA criteria. These Pfams correspond to eukaryote-specific families that did not duplicate.

#### Sister group identification

For each eukaryotic clade in trees also containing prokaryotic sequences the sister group was identified in an unrooted mode. By doing so, the eukaryotic clade initially had two candidate sister groups. Eukaryotic sequences in a sister group, if present, were ignored, as they could reflect HGT events, contaminations, tree artefacts or true additional acquisitions. To infer the actual sister group it was first checked if one of the two candidate sister groups was more likely by checking if one of them consisted only of Asgard archaea, TACK archaea, Asgard and TACK archaea, alphaproteobacteria, beta/gammaproteobacteria, or alpha/beta/gammaproteobacteria. If so, that clade was chosen as the actual sister group. If both sister groups had the same identity or if both groups had another identity than the ones described above, the tree was rooted on the farthest leaf from the eukaryotic clade. In many cases the last common ancestor of the taxa in the sister group was Bacteria, Archaea or cellular organisms (“LUCA”) according to the NCBI taxonomy. Such wide taxonomic assignments likely reflect extensive horizontal gene transfers among distantly related prokaryotes. In these cases it was checked if one of the previously mentioned groups or otherwise a particular phylum or proteobacterial class comprised a majority of the prokaryotic taxa to get a more precise sister group classification.

We observed that in a substantial number of cases there was another eukaryotic clade with LECA nodes in the sister group of a eukaryotic clade. These cases could reflect a duplication and subsequent loss in prokaryotes but probably reflect tree artefacts. Therefore these clades were ignored for the branch length analysis. Eukaryotic clades with LECA nodes that were nested, i.e. they had exactly the same prokaryotic sister group, were merged.

#### Branch length analysis

Multiple branch lengths were calculated in clades containing LECA nodes. For the stem length (sl) the distance to the acquisition node – the node uniting the eukaryotic clade and its prokaryotic sister – was calculated for each LECA node. This distance was divided by the median of the distances from the LECA node to the eukaryotic leaves (eukaryotic branch lengths (ebl)) to correct for rate differences between orthologous groups as done before^9^. In case of multiple possible paths due to duplications, the minimum of these distances was used as the sl, since it was closest to sl values from zero-duplication clades (Extended Data Fig. 5). To calculate the duplication length (dl) a similar approach was followed, using the duplication node instead of the acquisition node.

### Comparison between Pfam trees based on two and five groups subsampling

As mentioned before, the eukaryotic sequences that were in BBHs between five supergroups or between Opimoda and Diphoda species were selected for tree inference in the Pfam analysis. The two collections of annotated trees were compared. Many more duplication and especially LECA nodes had a higher support value in the Opimoda-Diphoda BBHs trees (Extended Data Fig. 6). Furthermore, the number of unclassified eukaryotic nodes was substantially higher in the between-supergroup BBHs trees (2,819 and 1,716, respectively). Therefore, we decided to show our findings on the Opimoda-Diphoda BBHs trees.

### Combining eukaryote-only Pfam families with prokaryotic donations in their clan

The classification of protein families into Pfams is not based on their taxonomic level. A Pfam present only in eukaryotes can therefore be the result of a duplication event instead of a bona fide invention. To distinguish these possible scenarios we used the Pfam clans, in which related Pfam families are combined. If there were only eukaryote-only Pfams in a clan based on our analysis, these Pfams were merged into one invention event. If there was only one Pfam with an acquisition form prokaryotes and for this Pfam there was only one acquisition, the eukaryote-only Pfams were combined with this acquisition. If there were multiple acquisitions in a clan, a profile-profile search with HH-suite3 v3.0.3^47^ was performed to assign eukaryote-only Pfams to an acquisition. Per acquisition in a clan an alignment was made from the tree sequences in the corresponding eukaryotic clade with MAFFT L-INS-i v7.310^39^. Profile HMMs were made of these alignments (hhmake –M 50) and they were combined in a database (ffindex_build). The eukaryote-only Pfam HHMs were searched against the acquisition HHM database per clan with hhsearch. Each Pfam was assigned to the acquisition that had the best score.

### Functional annotation

Functional annotation of sequences was performed using emapper-1.0.3^48^ based on eggNOG orthology data^32^. Sequence searches were performed using DIAMOND v0.8.22.84^49^.

The most common KOG functional category among the tree sequences of a LECA node was chosen as the function of the LECA node. If there was not one function most common, the node was annotated as S (function unknown). For the functional annotation of duplication nodes a Dollo parsimony approach was used. For this we checked if there was one single annotation shared between LECA nodes at both sides, ignoring unknown functions. If this was not the case but the parent duplication node (if present) had a function, this function was also used for the focal duplication node. In the figures the names of the categories were shortened for increased readability: Translation/J (Translation, ribosomal structure and biogenesis), RNA processing/A (RNA processing and modification), Transcription/K, Replication/L (Replication, recombination and repair), Chromatin/B (Chromatin structure and dynamics), Cell cycle/D (Cell cycle control, cell division, chromosome partitioning), Nuclear structure/Y, Defense mechanisms/V, Signal transduction/T (Signal transduction mechanisms), Cell wall/membrane/M (Cell wall/membrane/envelope biogenesis), Cell motility/N, Cytoskeleton/Z, Extracellular structures/W, Intracellular trafficking/U (Intracellular trafficking, secretion, and vesicular transport), Protein modification/O (Posttranslational modification, protein turnover, chaperones), Energy/C (Energy production and conversion), Carbohydrates/G (Carbohydrate transport and metabolism), Amino acids/E (Amino acid transport and metabolism), Nucleotides/F (Nucleotide transport and metabolism), Coenzymes/H (Coenzyme transport and metabolism), Lipids/I (Lipid transport and metabolism), Inorganic ions/P (Inorganic ion transport and metabolism), Secondary metabolites/Q (Secondary metabolites biosynthesis, transport and catabolism), General function prediction only/R, Function unknown/S.

The same approach was used to assign cellular components to LECA and duplication nodes, using a custom set of gene ontology terms: extracellular region (GO:0005576), cell wall (GO:0005618), cytosol (GO:0005829), cytoskeleton (GO:0005856), mitochondrion (GO:0005739), cilium (GO:0005929), plasma membrane (GO:0005886), endosome (GO:0005768), vacuole (GO:0005773), peroxisome (GO:0005777), cytoplasmic vesicle (GO:0031410), Golgi apparatus (GO:0005794), endoplasmic reticulum (GO:0005783), nuclear envelope (GO:0005635), nucleoplasm (GO:0005654), nuclear chromosome (GO:0000228) and nucleolus (GO:0005730).

### Predicting the number of genes in LECA

We used a linear regression model to predict the number of genes in LECA based on the inferred number of Pfam domains in LECA. For this, we used the number of sufficiently long Pfam domains (see ‘Pfam assignment’ above) and the number of protein-coding genes in the eukaryotes in our dataset. The assumptions of a normal distribution of gene values at each Pfam domain value and equal variance were reasonably met after log transformation.

### Statistical analysis

Overrepresentations of functions and localisations in duplications, inventions and innovations, and overrepresentations of sister groups in duplications and duplication tendencies were tested by comparing odds ratios with Fisher’s exact tests (only pairwise comparisons of functions for inventions and localisations for innovations due to small sample sizes) or χ^2^ contingency table tests (rest). Differences in branch lengths were assessed with a Kruskal-Wallis test, followed by Mann-Whitney *U* tests upon a significant outcome of the Kruskal-Wallis test, which was always the case. All performed tests were two-sided. In all cases of multiple comparisons, the *p*-values were adjusted to control the false discovery rate.

## Data availability

The phylogenetic trees and their annotations are available in figshare with the identifier doi:10.6084/m9.figshare.10069985. Additional data are available from the corresponding authors upon request.

## Code availability

The code used to annotate the phylogenetic trees can be accessed in Github (https://github.com/JulianVosseberg/feca2leca).

## Extended Data

**Extended Data Fig. 1.**
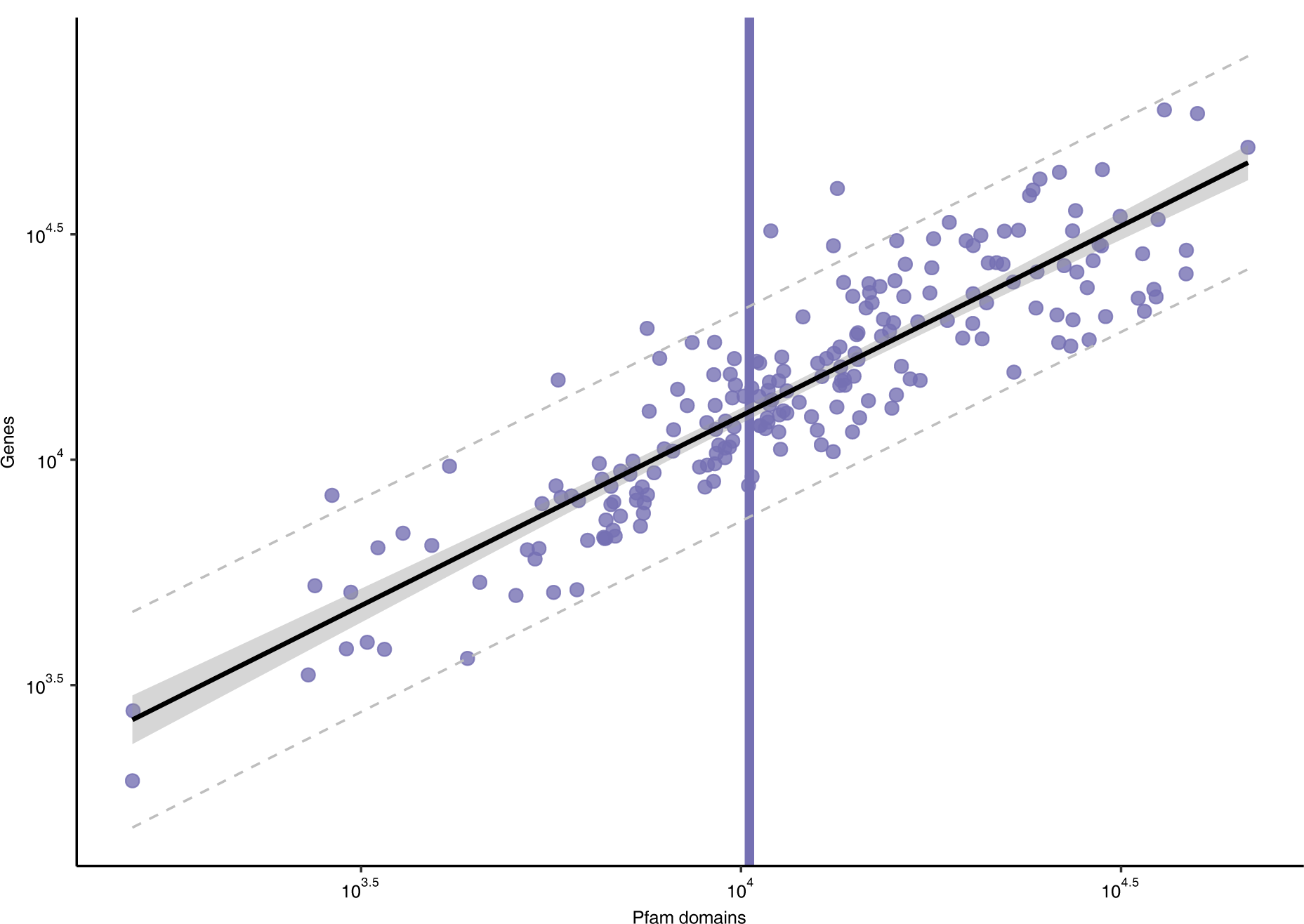
Estimating the number of LECA genes from the number of Pfam domains with linear regression. Scatter plot showing the number of Pfam domains and protein-coding genes in present-day eukaryotes, with each dot representing one genome. The regression line (black) and its 95% confidence (filled grey) and prediction intervals (dashed grey) are depicted. The vertical line corresponds to the obtained number of LECA Pfam domains.

**Extended Data Table 1.** Comparison of different datasets.

|  | KOG-to-COG | Pfam Opimoda-Diphoda BBHs | Pfam between-supergroup BBHs |
| --- | --- | --- | --- |
| Acquisitions | 3,460 | 4,340 | 4,003 |
| Inventions | 883 | 1,336 | 1,347 |
| Duplications | 4,888 | 4,583 | 4,626 |
| LECA families | 9,231 | 10,259 | 9,976 |
| Multiplication factor | 2.12 | 1.81 | 1.86 |

**Extended Data Table 2.** Most expanded acquisitions or inventions during eukaryogenesis.

| Pfam | Ancestry | Number of LECA families |
| --- | --- | --- |
| **Total** |  |  |
| Mitochondrial carrier* | Invention | 123 |
| Protein kinase | Planctomycetes | 106 |
| RING-finger/U-box | Asgard archaea | 93 |
| PH domain | *Haloplasma* | 82 |
| Ubiquitin | Asgard archaea | 74 |
| C2 domain | Prokaryotes | 72 |
| RNA recognition motif | $\alpha\beta\gamma$-proteobacteria | 71 |
| POZ domain | Chlamydiae | 50 |
| FYVE/PHD zinc finger | Invention | 46 |
| Tetratricopeptide repeat | Spirochaetes | 42 |
| **Asgard archaea** |  |  |
| RING-finger/U-box | Asgard archaea | 93 |
| Ubiquitin | Asgard archaea | 74 |
| Ubiquitin-conjugating enzyme | Asgard archaea | 37 |
| Ras family (Rab) | Asgard archaea | 25 |
| Vps51 domain superfamily | Asgard archaea | 19 |
| Cyclin | Asgard archaea | 18 |
| Helix-turn-helix | Asgard archaea | 16 |
| Leucine-rich repeat | Asgard archaea | 15 |
| ABC transporter | Asgard archaea | 14 |
| **Alphaproteobacteria** |  |  |
| Sterile alpha motif | Alphaproteobacteria | 10 |
| Galactosyltransferase | Alphaproteobacteria | 9 |
| EF-hand 8 | Alphaproteobacteria | 8 |
| Iron/zinc purple acid phosphatase-like protein C | Alphaproteobacteria | 5 |
| DDE superfamily endonuclease | Alphaproteobacteria | 5 |
| Acyl-CoA oxidase/dehydrogenase N-terminal | Alphaproteobacteria | 4 |
| GTP binding protein | Alphaproteobacteria | 4 |
| Ferric reductase | Alphaproteobacteria | 4 |

\*A mitochondrial carrier protein typically contains three of these domains.

**Extended Data Fig. 2.**
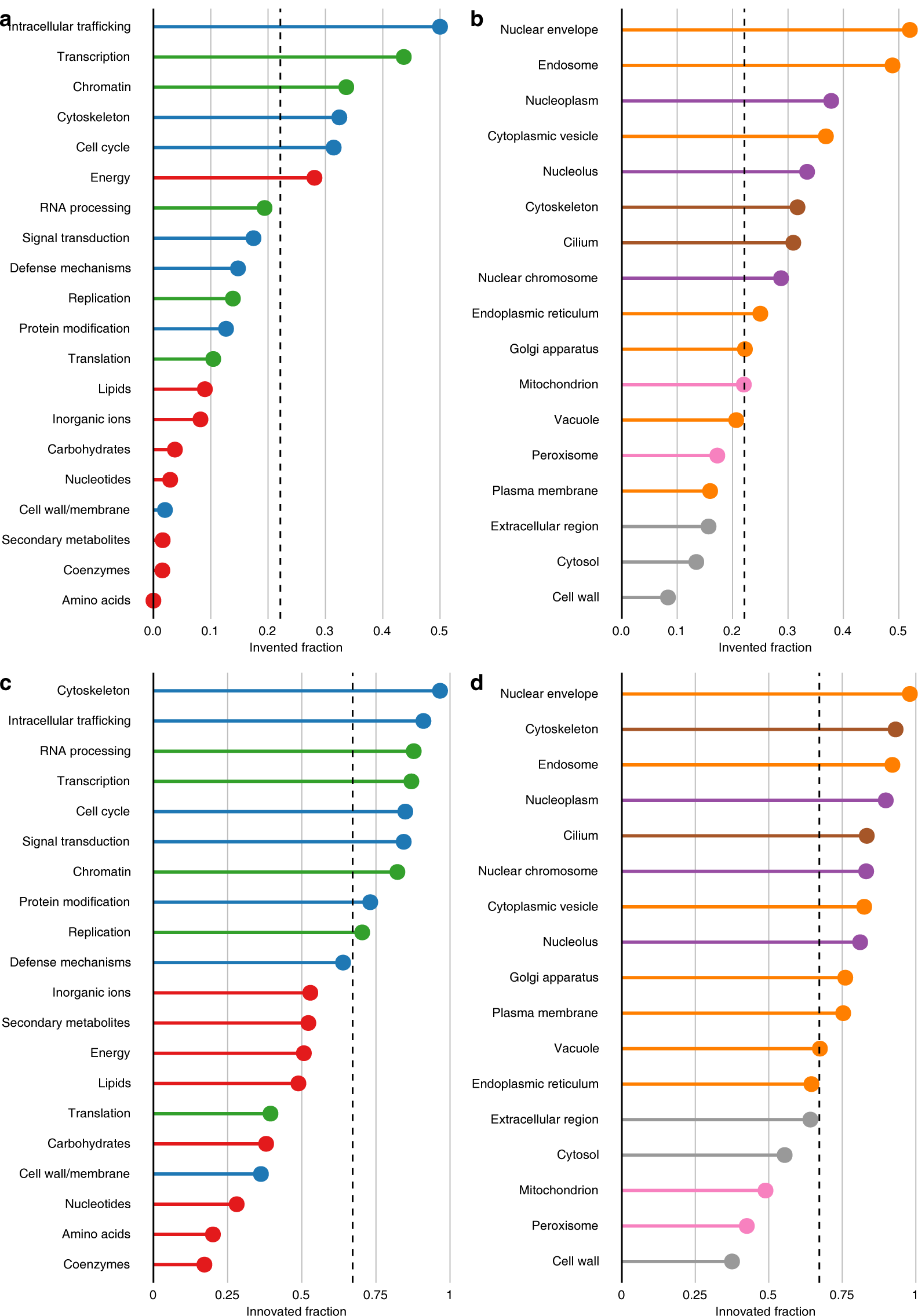
Fraction of LECA families resulting from inventions. **a**, Contribution of inventions to LECA families performing different functions. 81% of pairwise comparisons were significantly different (Supplementary Fig. 3). **b**, Contribution of inventions to LECA families performing their function in different cellular components. 51% of pairwise comparisons were significantly different (Supplementary Fig. 4). **c**, Fraction of LECA families resulting from either an invention or duplication – a eukaryotic innovation – according to functional category. 85% of pairwise comparisons were significantly different (Supplementary Fig. 5). **d**, Fraction of LECA families resulting from an innovation according to cellular localisation. 75% of pairwise comparisons were significantly different (Supplementary Fig. 6). **a-d**, Dashed lines indicate the overall invented or innovated fraction.

**Extended Data Fig. 3.**
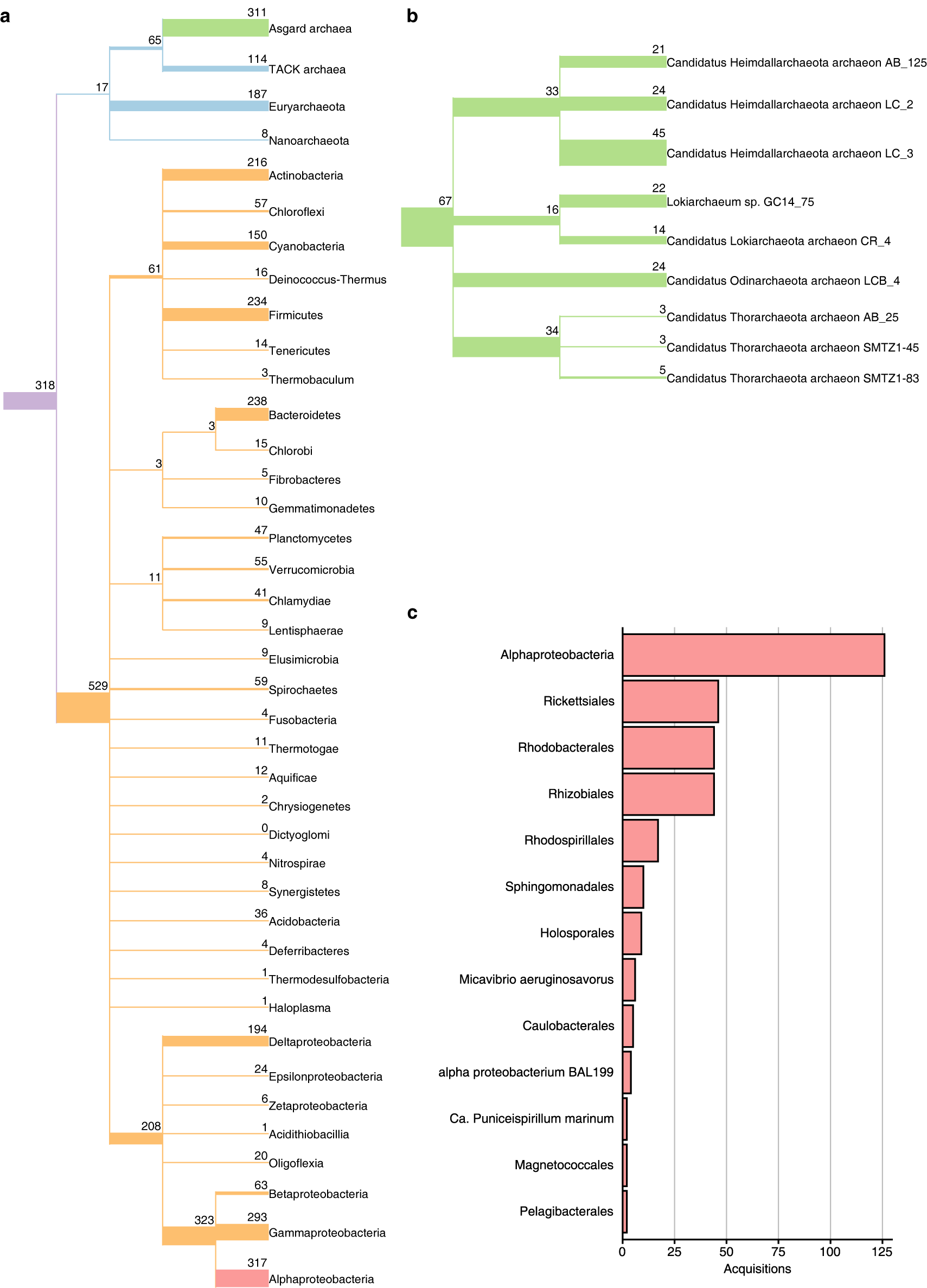
Phylogenetic origin of acquired Pfams. **a, b**, Phylogeny of the prokaryotes (**a**) and Asgard archaea (**b**) present in our dataset based on the NCBI taxonomy. The branch widths and numbers indicate the number of acquisitions from a group. **c**, Number of acquisitions from different alphaproteobacterial orders or a combination of multiple orders (‘Alphaproteobacteria’).

**Extended Data Fig. 4.**
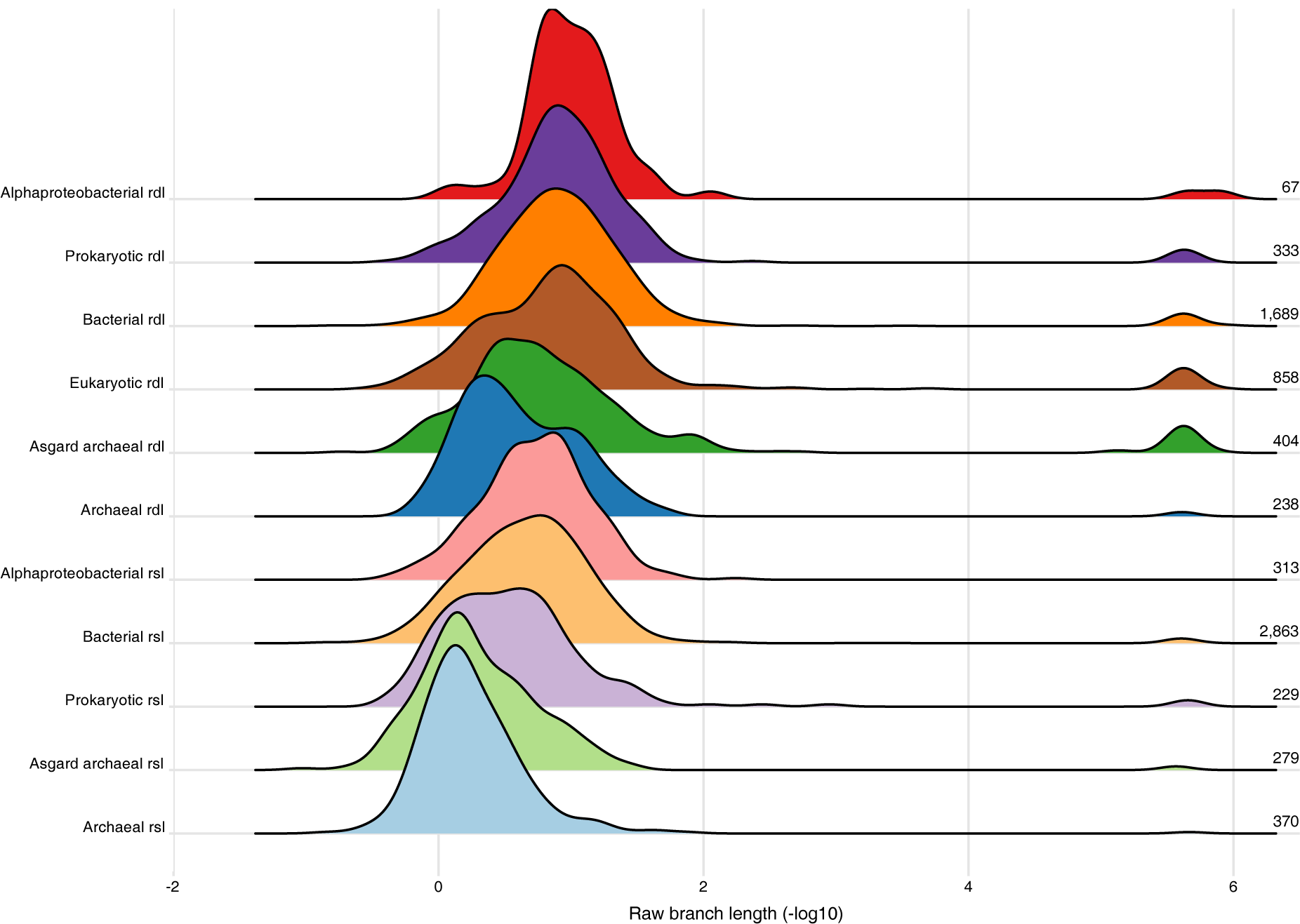
Effect of branch length normalisation. Ridgeline plot showing the distribution of uncorrected stem (rsl) or duplication lengths (rdl), depicted as the additive inverse of the log-transformed values. Numbers indicate the number of acquisitions or duplications for which the branch lengths were included. The low peaks at very short branch lengths are an artefact from near-zero branch lengths. Groups are ordered based on the median value of rsls and rdls.

**Extended Data Fig. 5.**
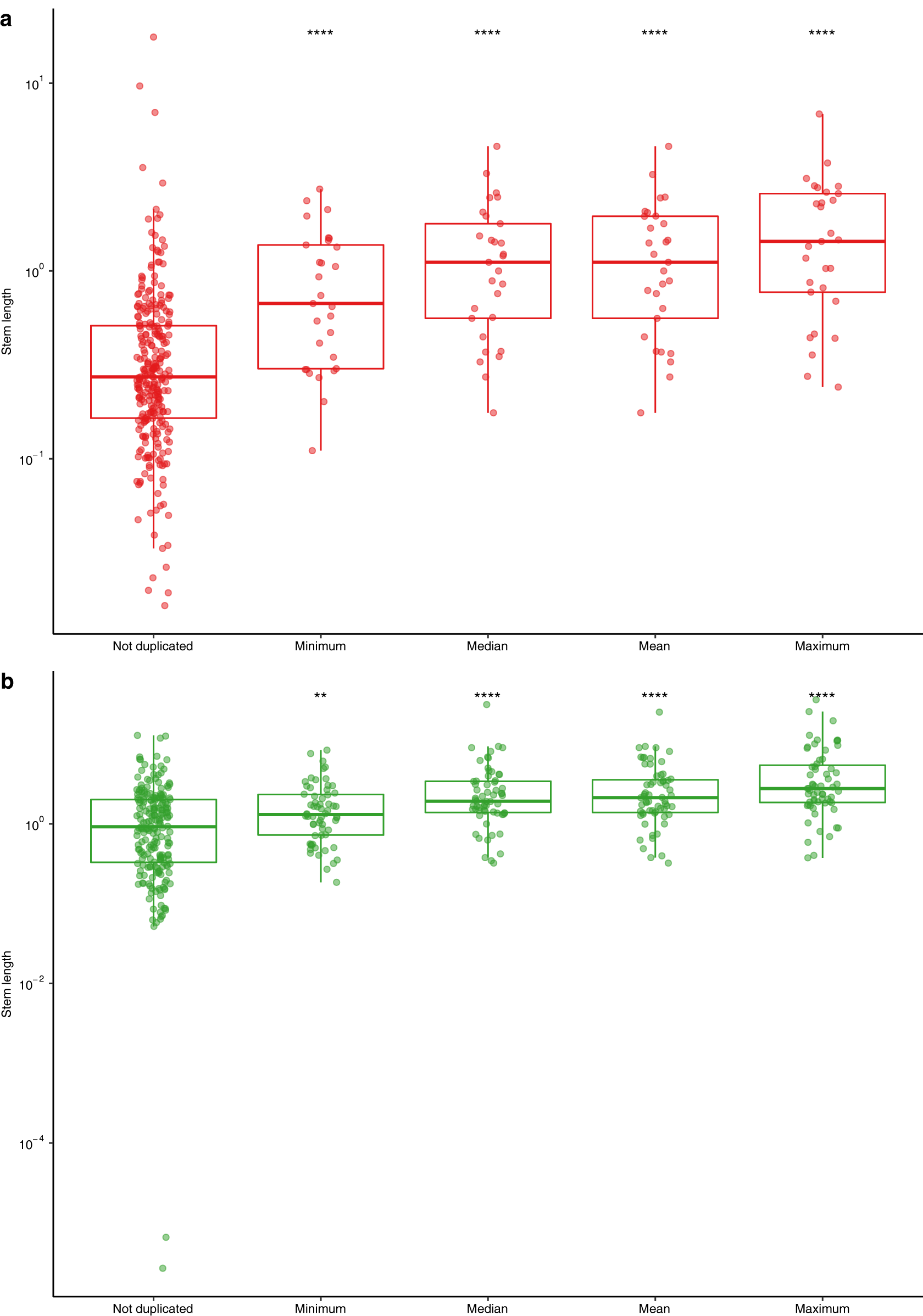
Testing effect of using different measures to calculate the stem lengths in case of duplications. **a, b**, Distribution of alphaproteobacterial (**a**) and Asgard archaeal (**b**) stem lengths. Alphaproteobacterial stem lengths with Magnetococcales as sister group were removed. Asterisks indicate the significance levels of Mann-Whitney *U*-tests with the ‘Not duplicated’ group as reference group (****: P < 0.0001; **: P < 0.01).

**Extended Data Fig. 6.**
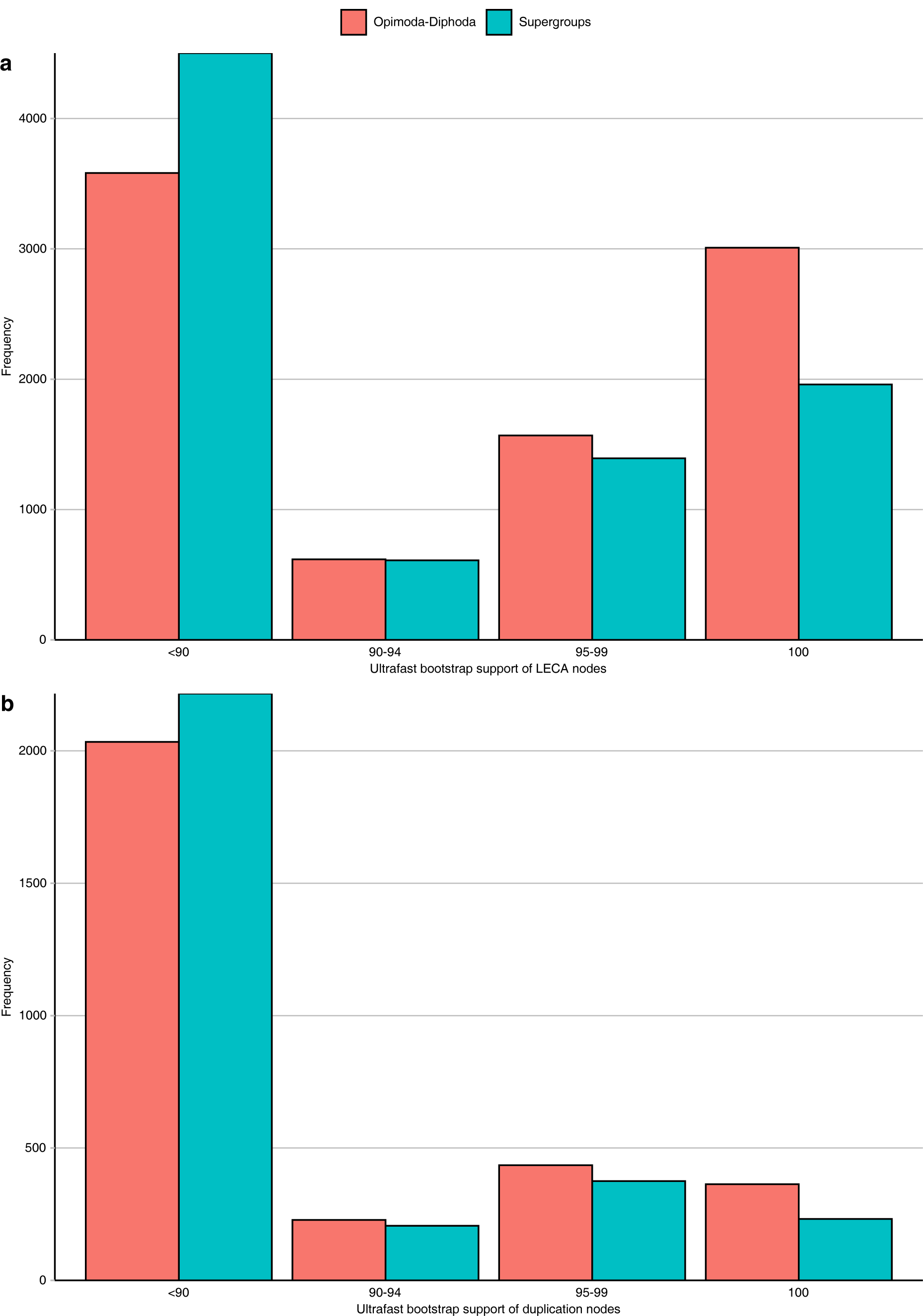
Comparison of the Opimoda-Diphoda and between-supergroup BBHs subsampled trees. **a, b**, Number of LECA (**a**) or duplication (**b**) nodes with a particular ultrafast bootstrap support value.

## Notes

### Competing Interest Statement

The authors have declared no competing interest.

