## Supplementary Information for "Timing the origin of eukaryotic cellular complexity with ancient duplications"

**Table of contents**

**1. Supplementary Discussion**
**2. Supplementary References**
**3. Supplementary Figures**

**Supplementary Discussion**

***Data sets used***

We tested two different data sets. The KOG-to-COG gene family clusters[10] are a set specifically constructed to study duplications during eukaryogenesis and were therefore an ideal starting point. To get an even more complete picture of duplications we decided to use the Pfam database. By using this database we circumvented the need to use orthologous groups or infer homology. For certain families the Pfam domains correspond to full-length genes, whereas for others it is only a domain or even a motif. Although certain domain duplications are not fully independent of each other due to their presence in a single gene upon duplication, it is not unlikely that truly separated genes co-duplicated as well. Ideally, one would want to define the unit, either a domain or full-length gene, that is best suited for the timescale the evolution of a particular gene is studied, although this might be even different for the acquisition/invention level and the LECA level. This is especially probable given the abundance of gene fusion events during eukaryogenesis[50].

***Sister group identity***

7% of the acquisitions had an unclear prokaryotic ancestry. Both bacteria and archaea were present in the sister group with no phylum comprising a majority. A tentative explanation is that the identity of the donor is obscured due to post-acquisition HGT among distantly related prokaryotes. The tendency of these acquisitions to duplicate was similar to the Pfams with an archaeal ancestry (Fig. 2a, Supplementary Fig. 7a). Likewise, the multiplication factor was high, being second only to the Asgard archaeal group (Fig. 2b, Supplementary Fig. 7b). This suggests that a large fraction of this group reflect genes present in the host lineage. Furthermore, a relatively large fraction of these acquisitions had another eukaryotic clade with LECA families in their sister group (34%, between 3 and 10% for the other groups), indicating that some of these acquisitions are placed in an incorrect, deep phylogenetic position.

***Branch lengths analysis***

Since a duplication between the acquisition and LECA resulted in multiple LECA nodes, there are multiple estimates for the stem length in case of duplications. The stem lengths of acquisitions that happened simultaneously should approximate the same value. Assuming the deep mitochondrial origin outside the alphaproteobacteria[7], all acquisitions with alphaproteobacteria as sister group should correspond to the same event, namely the divergence of the pre-mitochondrial and alphaproteobacterial lineages. We tested multiple measures for those acquisitions that had subsequent duplications and observed that the shortest length was most consistent with the branch lengths obtained from non-duplicated acquisitions (Extended Data Fig. 5a). It should be noted that even using the shortest could not fully account for the difference in stem lengths. We also tested multiple measures for acquisitions with an Asgard archaeal sister group, although the exact phylogenetic position of eukaryotes within Asgard archaea has not yet been resolved[51]. Also for these acquisitions the shortest length was the best option (Extended Data Fig. 5b). Therefore, we used the shortest branch length to time acquisitions and duplications.

This indicates that in most cases there was an accelerated evolutionary rate in at least one of the paralogues, which could not be (fully) corrected for by the post-LECA branch lengths. These gene families may have followed a pattern of episodic, rapid sequence evolution during eukaryogenesis followed by slower, conservative evolution after LECA in the different eukaryotic lineages. Such a pattern would fit into a 'Big Bang' hypothesis for eukaryogenesis[52] and may underlie the many functional innovations during the prokaryote-to-eukaryote transition, compared to more recent, post-LECA evolution.

We also tested the effect of the normalisation using the post-LECA branches and observed that the normalisation mainly affected the Asgard archaeal and invention duplication lengths (Extended Data Fig. 4). For both, normalisation increased the number of duplications predating mitochondrial endosymbiosis, making the mitochondrial acquisition a slightly later event. Overall, the observed patterns were independent of the normalisation.

A limitation of stem lengths is that inferred acquisitions are the result of taxon sampling (i.e. which of the present-day organisms have been discovered, sequenced and/or included in the analysis) and historical contingency (i.e. which lineages have not gone extinct). They represent the earliest possibility of the actual acquisition. Duplication nodes, on the other hand, very likely reflect true biological events. Moreover, they represent the latest possibility of the actual acquisition. Including duplications in branch length analyses to time key events therefore not only provides insights into the successive stages of eukaryogenesis, but also gives better estimates for the timing of the acquisition of genes from prokaryotes.

***Comparison with Tria* et al.[17]**

Our conclusions are in stark contrast with a recent preprint[17]. However, as mentioned in the main text, that study was unable to recover greatly expanded protein families. The family that according to this preprint was most duplicated during eukaryogenesis was the dynein light chain family with 12 duplications. They did not recover well-documented more expanded protein families such as protein kinases and small GTPases[11,13], which we were able to recover (Extended Data Table 2). In general, they inferred very few duplications, which stands in stark contrast with our study and Makarova *et al.* (2005)[10]. In addition, as the analysis by Tria *et al.* is based on phylogenies containing only eukaryotic sequences, it cannot possibly differentiate duplications that occurred in the LECA stem from those predating its divergence from prokaryotic relatives. Finally, they did not include the Asgard archaea in their analysis, which are crucial for any inference about eukaryogenesis. This might explain how the duplications in the cytoskeletal and ubiquitin systems were not correctly identified as duplications in archaeal acquisitions[4,5] in their analysis.

### Supplementary Figures

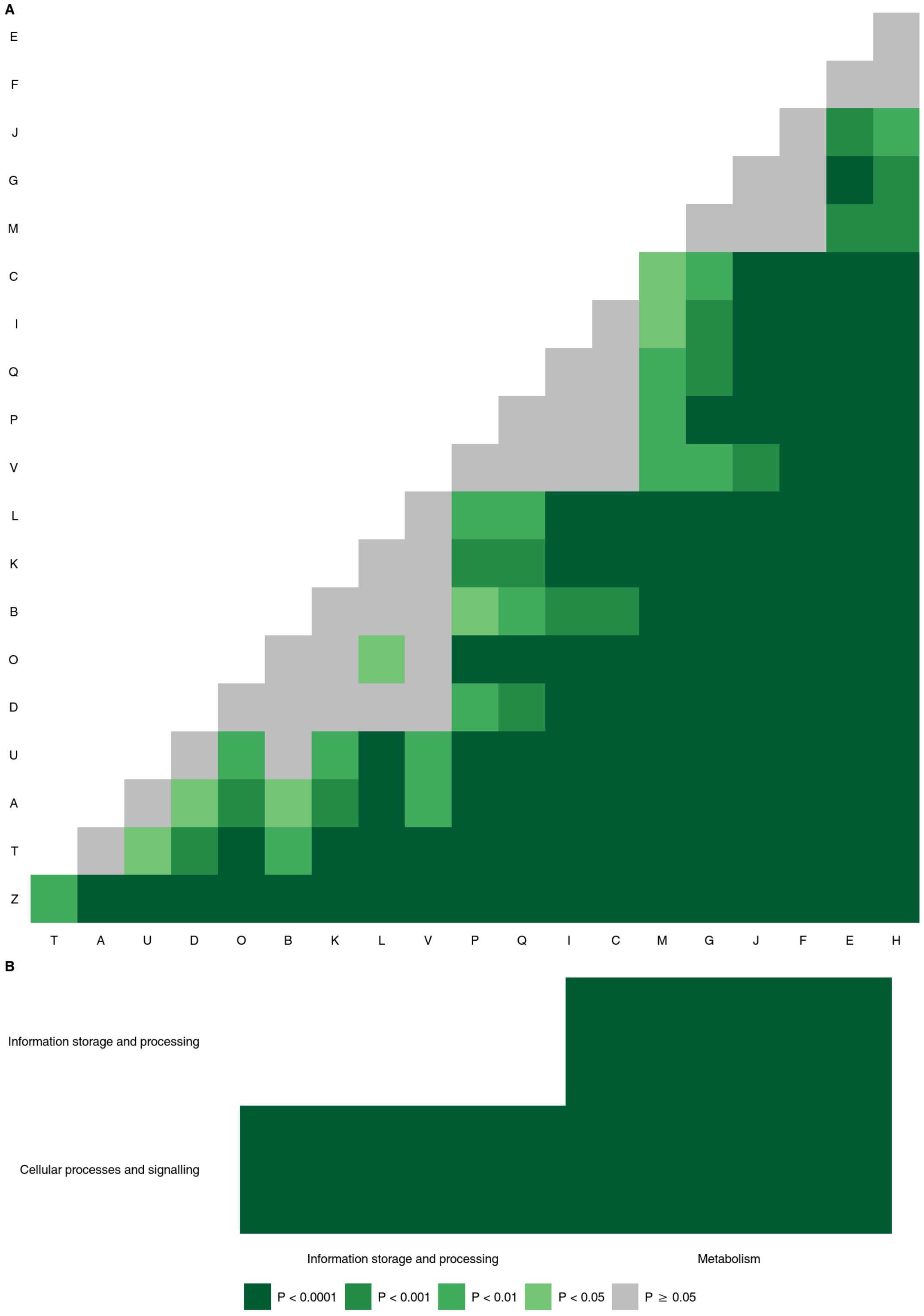


**Supplementary Fig. 1 | Contribution of duplications to families with a particular function.**

Pairwise comparisons of different functional categories (**A**) and the corresponding broad categories (**B**). For the meaning of the abbreviations, see ‘Methods’.

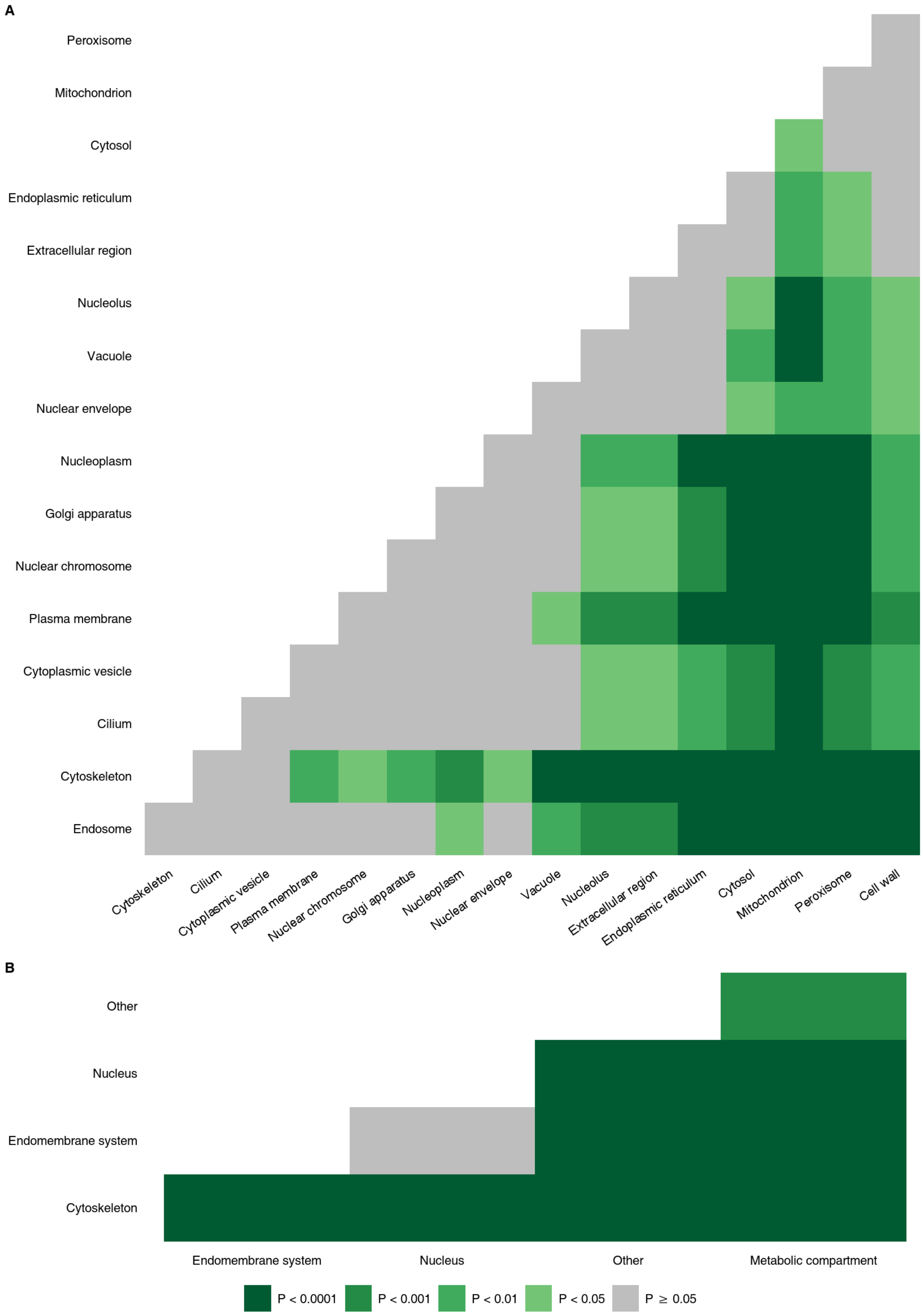


**Supplementary Fig. 2 | Contribution of duplications to families with a particular cellular localisation.**

Pairwise comparisons of different localisations (**A**) and the corresponding broad categories (**B**).

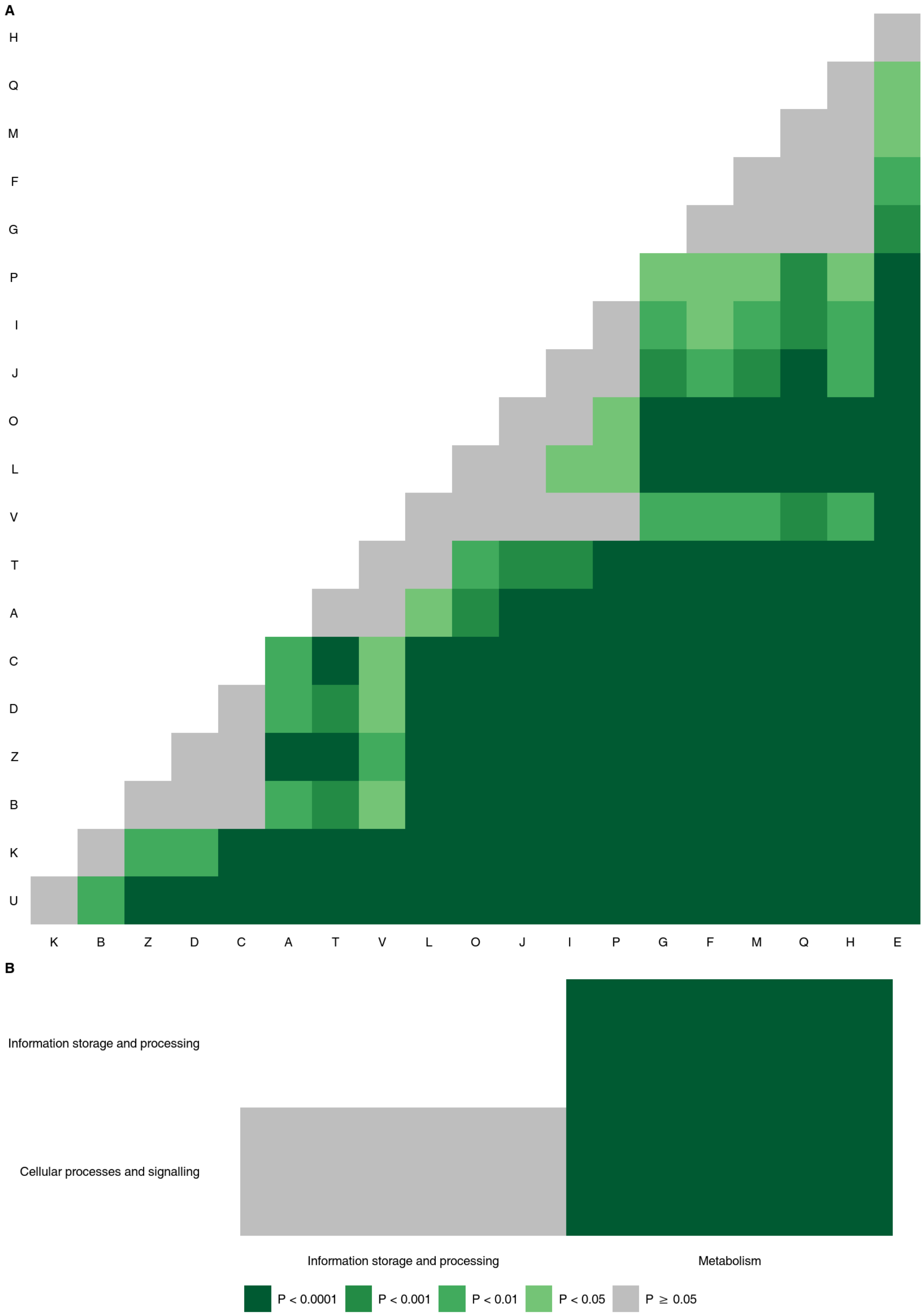


**Supplementary Fig. 3 | Contribution of inventions to families with a particular function.** Pairwise comparisons of different functional categories (**A**) and the corresponding broad categories (**B**).

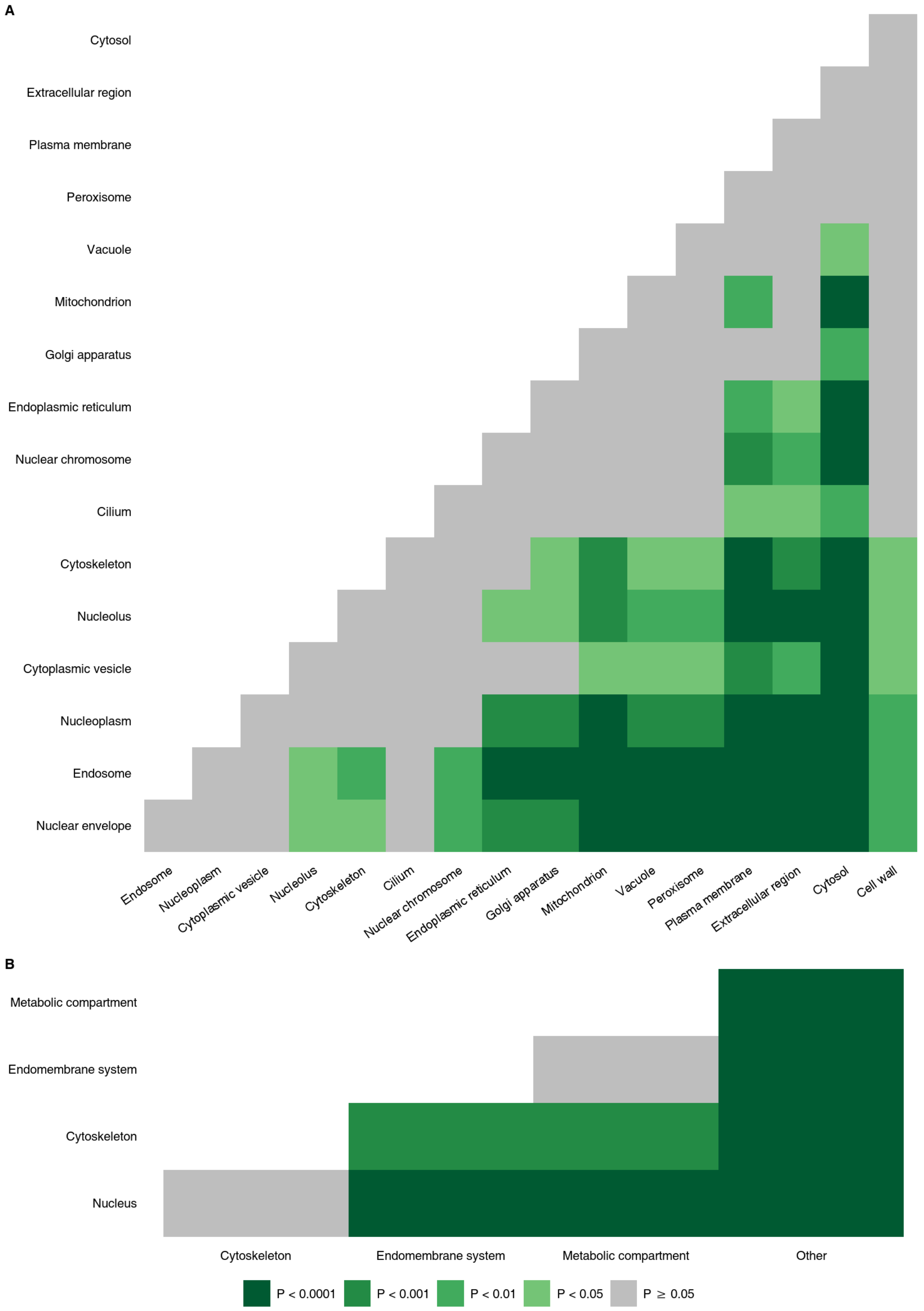


**Supplementary Fig. 4 | Contribution of inventions to families with a particular cellular localisation.**

Pairwise comparisons of different localisations (**A**) and the corresponding broad categories (**B**).

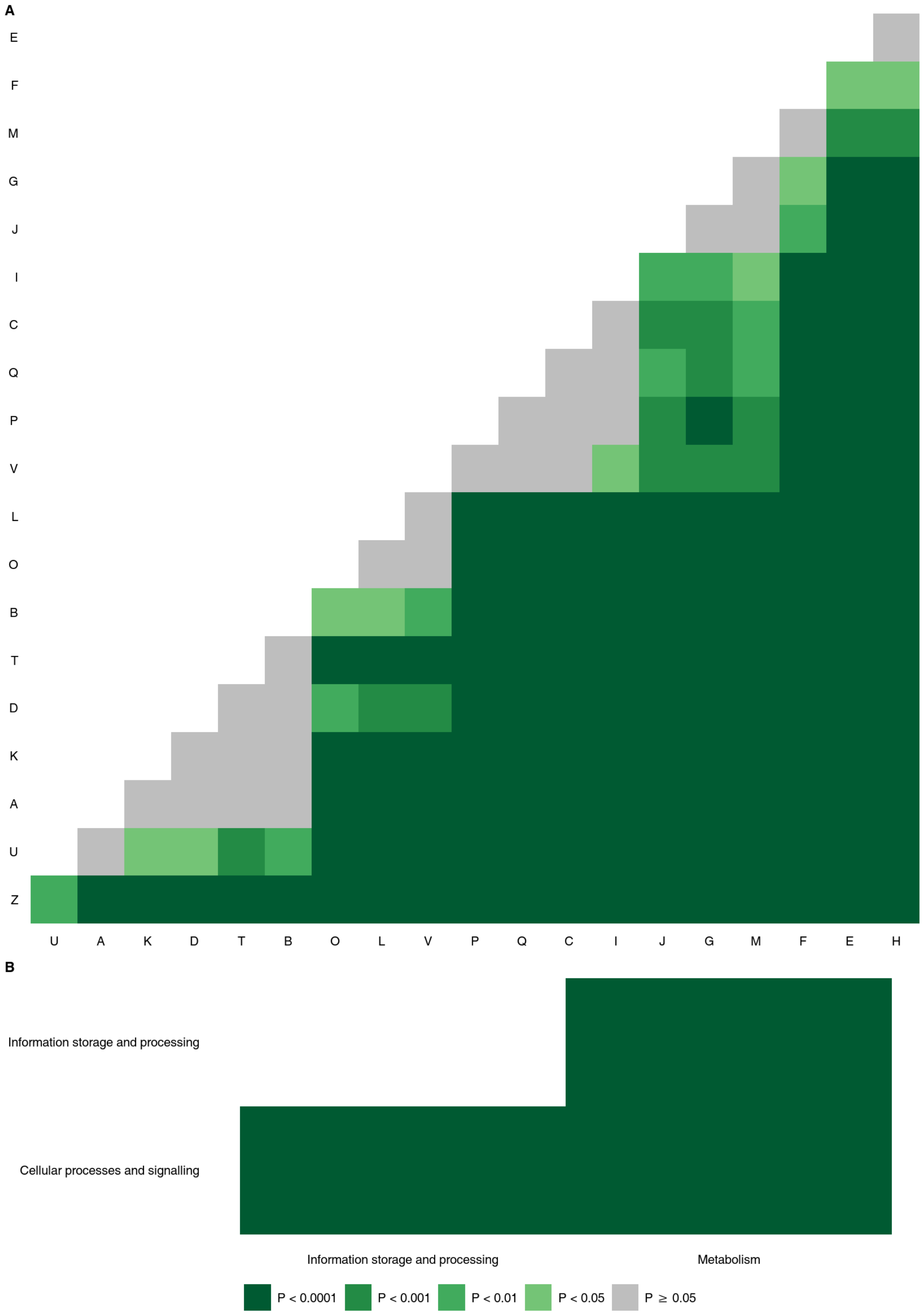


**Supplementary Fig. 5 | Contribution of innovations to families with a particular function.**

Pairwise comparisons of different functional categories (**A**) and the corresponding broad categories (**B**).

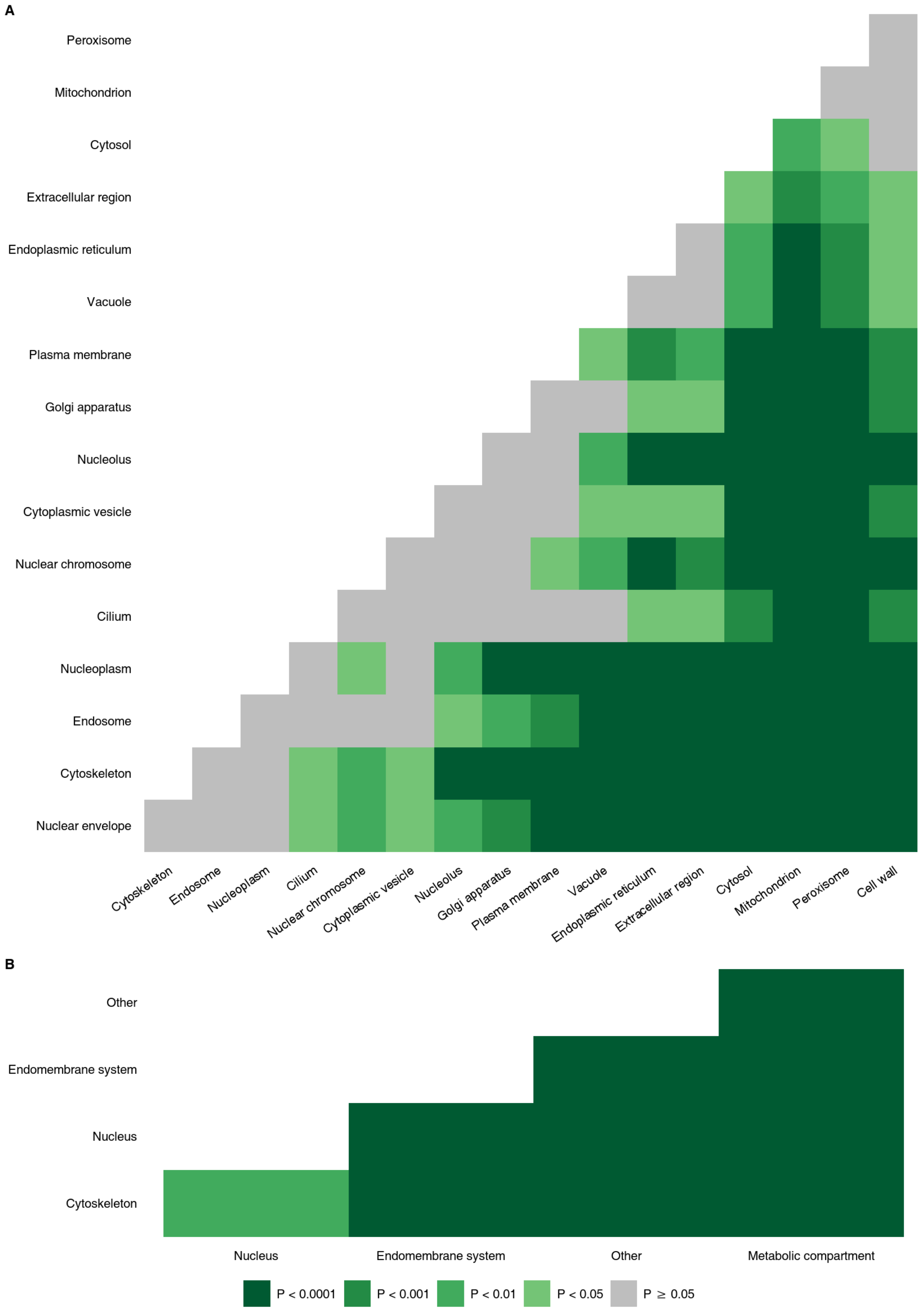


**Supplementary Fig. 6 | Contribution of innovations to families with a particular cellular localisation.**

Pairwise comparisons of different cellular localisations (**A**) and the corresponding broad categories (**B**).

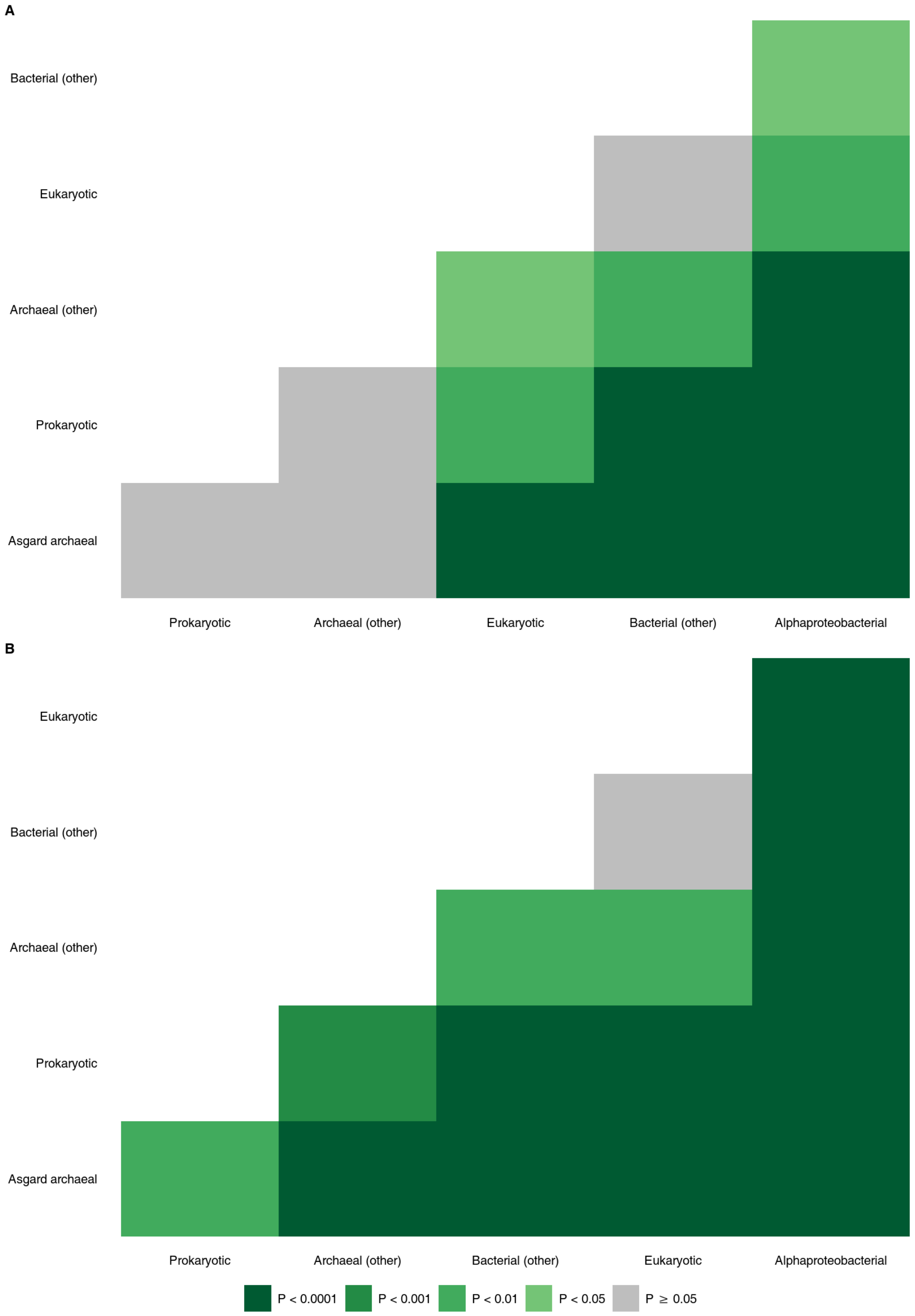


**Supplementary Fig. 7 | Contribution of different phylogenetic origins to duplications.** Pairwise comparisons of duplication tendencies (**A**) and multiplication factors (**B**).

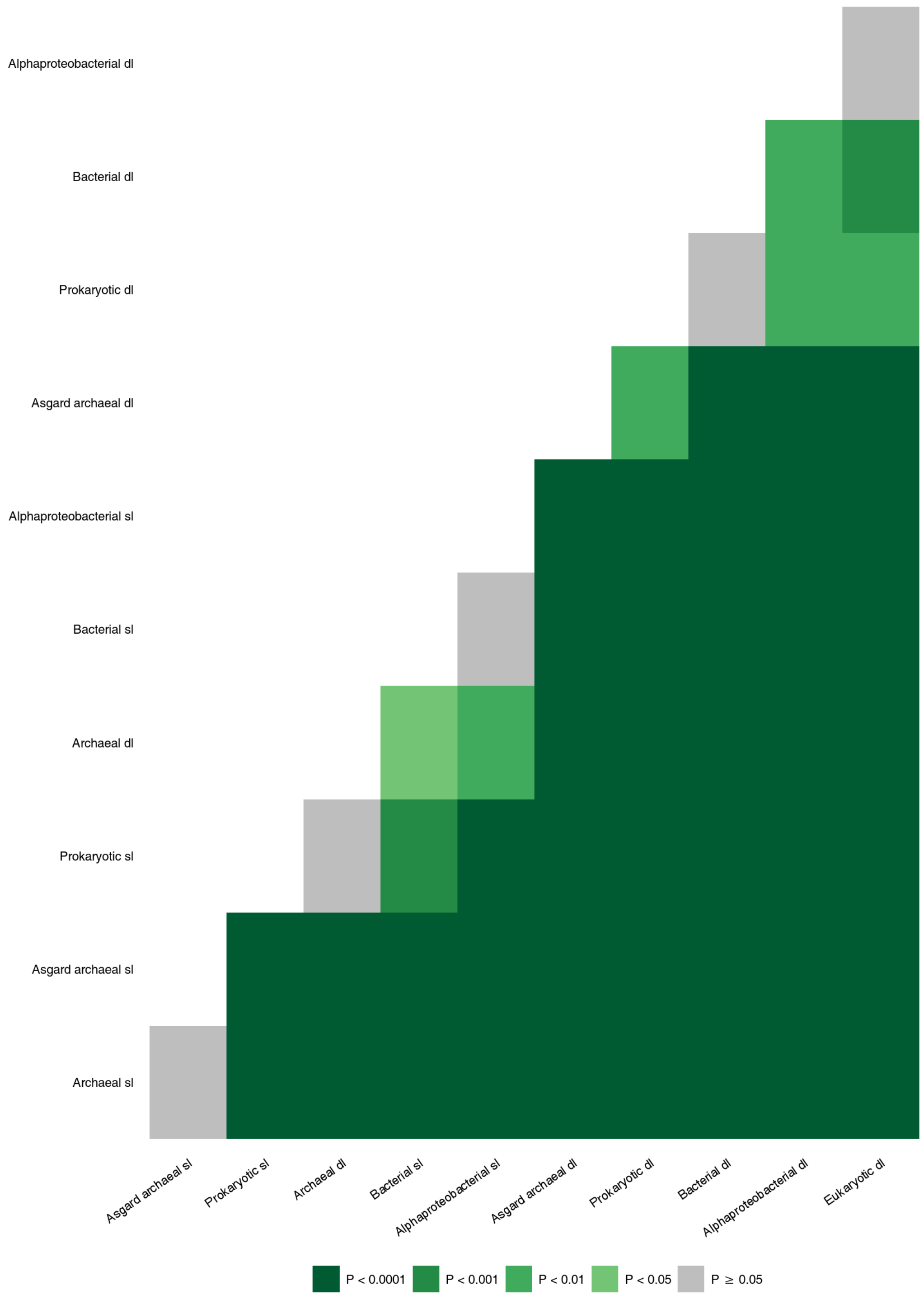


**Supplementary Fig. 8 | Comparison of branch lengths between different phylogenetic origins.**

sl: stem lengths; dl: duplication lengths.

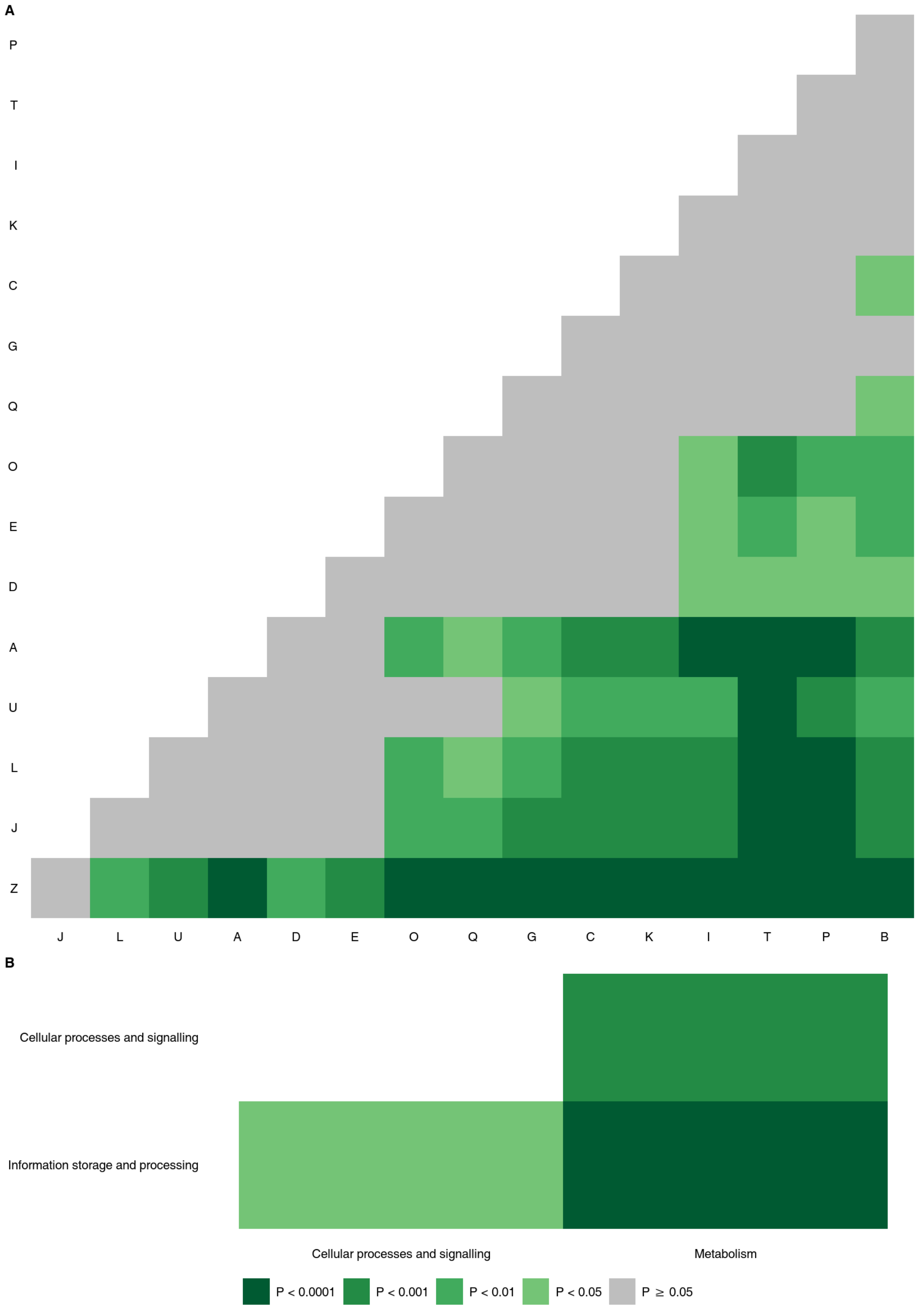


**Supplementary Fig. 9 | Comparison of duplication lengths between different functions.** Pairwise comparisons of different functional categories (**A**) and the corresponding broad categories (**B**).

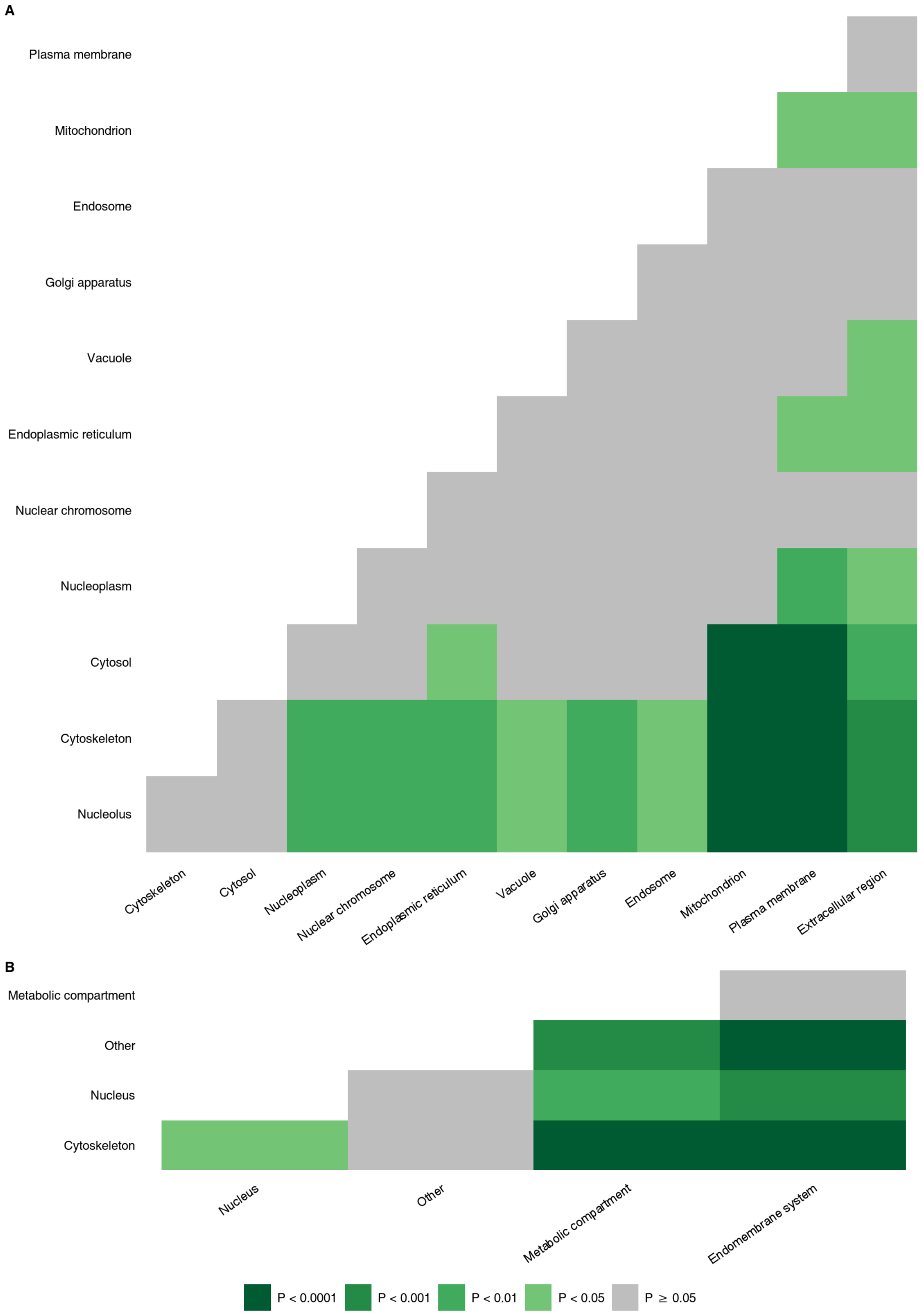


**Supplementary Fig. 10 | Comparison of duplication lengths between different cellular localisations.**

Pairwise comparisons of different cellular localisations (**A**) and the corresponding broad categories (**B**).
